# Cellular contractility coordinates cytoskeletal dynamics and cell behaviour during *Drosophila* abdominal morphogenesis

**DOI:** 10.1101/672006

**Authors:** Pau Pulido Companys, Anneliese Norris, Marcus Bischoff

## Abstract

During morphogenesis, cells undergo various behaviours, such as migration and constriction, which need to be coordinated. How this is achieved remains elusive. During morphogenesis of the *Drosophila* adult abdominal epidermis, larval epithelial cells (LECs) migrate directedly before constricting apically and undergoing apoptosis. Here, we study the mechanisms underlying the transition from migration to constriction. We show that LECs possess a pulsatile apical actomyosin network and that a change in network polarity underlies behavioural change. Exploring the properties of the contractile network, we find that the level of cell contractility impacts on the network’s behaviour, as well as on overall cytoskeletal architecture and cell behaviour. We also find that pulsed contractions occur only in cells with intermediate levels of contractility. Furthermore, increasing levels of the small Rho GTPase Rho1 disrupts pulsed contractions, and instead leading to cells that cycle between two states, characterised by a junctional cortical and an apicomedial actin network. Our results highlight that behavioural change relies on tightly controlled cellular contractility. Moreover, we show that constriction can occur without pulsed contractions, raising questions about their contribution to constriction.

## Introduction

During development, morphogenetic processes ultimately shape the organism (Lecuit and Le Goff, 2007). Such processes are driven by various cell behaviours, e.g. intercalation (Bertet et al., 2004; Irvine and Wieschaus, 1994), division (Gong et al., 2004), migration (Montell, 1999) and shape change (Butler et al., 2009), all of which need to be coordinated (Lecuit and Le Goff, 2007). Little is known about how this coordination is achieved, how cells switch behaviour and how different behaviours occur simultaneously.

Planar cell polarity (PCP) directs cell behaviour in the plane of the tissue, coordinating e.g. junctional remodelling (Bosveld et al., 2012; Zallen and Wieschaus, 2004), division orientation (Baena-Lopez et al., 2005; Mao et al., 2011) and migration (Keller, 2002). Migrating cells have a protruding front and a contracting back (Lauffenburger and Horwitz, 1996), further indicating cytoskeletal polarity. In contrast, apically constricting cells show radial cell polarity (RCP) (Mason et al., 2013).

Ultimately, cell behaviour depends on the cytoskeleton, particularly the actin cytoskeleton. Cell migration relies on protrusive activity, such as lamellipodia formation (Lauffenburger and Horwitz, 1996). Cell shape changes, including apical constriction, which reduces apical cell area, depend on actomyosin contractility (Barrett et al., 1997; Kinoshita et al., 2008; Nikolaidou and Barrett, 2004; Sawyer et al., 2010).

Increasing evidence suggests that the actin cytoskeleton shows dynamic rhythmic activity, such as actin flows during migration (Huang et al., 2013) and pulsed contractions during junctional remodelling, neuroblast ingression, apical constriction, basal constriction and vertebrate neural tube closure (Christodoulou and Skourides, 2015; He et al., 2010; Martin et al., 2009; Rauzi et al., 2010; Roh-Johnson et al., 2012; Simoes et al., 2017; Solon et al., 2009). Pulsed contractions are driven by periodic actomyosin contractions (Blanchard et al., 2010; Levayer and Lecuit, 2012; Martin et al., 2009; Mason and Martin, 2011). The ‘ratchet model’ describes cycles of pulsed contractions leading to cell area fluctuation, followed by stabilisation of the resulting smaller area (Martin et al., 2009).

Pulsed contractions are regulated by phosphorylation of Myosin II regulatory light chain (MRLC; Spaghetti squash (Sqh) in *Drosophila*) by Rho kinase (Rok) (Dawes-Hoang, 2005; Munjal et al., 2015; Vasquez et al., 2014), and its dephosphorylation by Myosin phosphatase (Fischer et al., 2014; Munjal et al., 2015; Valencia-Expósito et al., 2016). Upstream, the small GTPase Rho1 is involved in regulating actomyosin contractility in many contexts, from rear retraction in migrating cells (Ridley, 2003) to pulsed contractions (Mason et al., 2016; Munjal et al., 2015). During pulsed contractions, the activity of Rho1 is regulated by activating guanine nucleotide exchange factors (GEFs) and inhibitory GTPase-activating proteins (GAPs) (Graessl et al., 2017; Mason et al., 2016).

During formation of the adult abdominal epidermis of *Drosophila*, the larval epithelial cells (LECs) are replaced by the adult histoblasts (Bischoff and Cseresnyes, 2009; Madhavan and Madhavan, 1980; Ninov et al., 2007). We have shown previously that LECs undergo directed migration followed by a transition to apical constriction, which eventually leads to delamination and apoptosis (Bischoff, 2012) (Fig. 1A).

**Figure 1.**
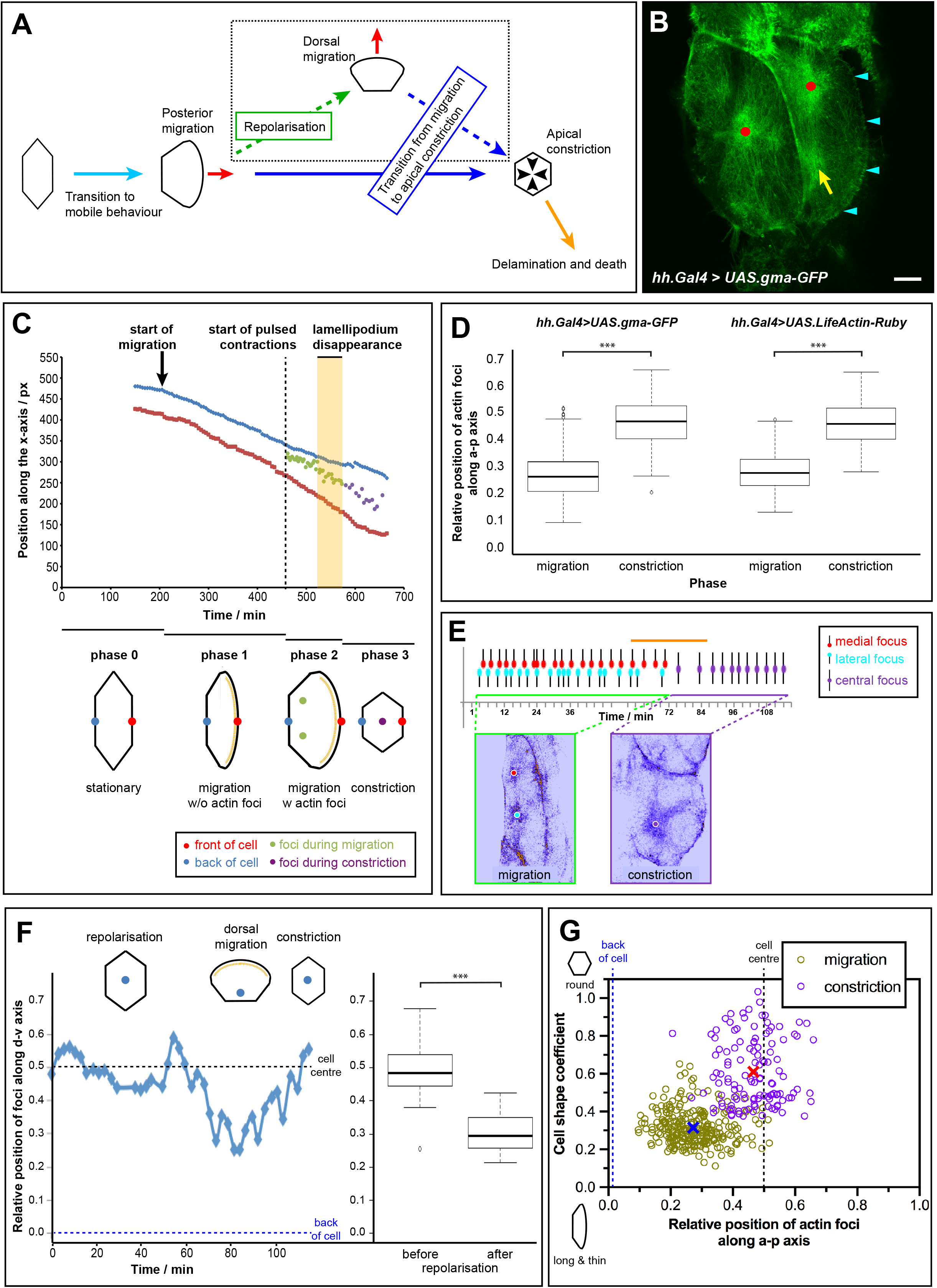
LECs undergo pulsed contractions that correlate with their behaviour. (**A**) LEC behaviour throughout morphogenesis (Bischoff, 2012). (**B**) Two LECs undergoing pulsed contractions. GMA-GFP labels F-actin. Left cell contracts; right cell migrates and contracts. Red dots, assembling actin foci; yellow arrow, disassembling actin focus; cyan arrowheads, lamellipodium. Note junctional actin pool. Bar, 10 μm. (**C**) LECs go through four phases of behaviour. Actin foci and front and back of cell were tracked over time. Foci are first positioned at back of cell, but once lamellipodium disappears (yellow area), foci move to cell centre. (**D**) Relative position of actin foci along a-p axis in GMA-GFP and LifeActin-Ruby pupae during migration and constriction. 0.5=cell centre, 0=cell back. (**E**) Periodicity of actin foci. Two foci alternate during migration; single focus during constriction. Orange line indicates period of lamellipodium disappearance; here, pulsing is less regular. (**F**) Actin foci during dorsal migration after repolarisation. Left: relative position of focus along d-v axis, tracked over time. Focus moves towards new cell back, as defined by newly formed lamellipodium. Right: after repolarisation, foci are positioned significantly more towards the ‘new’ cell back than before (p<0.001; n=3). (**G**) Cell shape *vs*. relative position of actin foci along a-p axis during migration and constriction. Cells that are long along d-v and thin along a-p axis tend to show foci in back, whereas round cells tend to have one focus in centre. Blue cross, 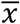(migration); red cross, 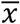(constriction). Anterior, left; dorsal, up.

To gain insights into the mechanisms underlying behavioural change, we studied the LEC cytoskeleton during the transition from migration to constriction. We show that LECs possess an apicomedial actomyosin network that undergoes pulsed contractions. This network is planar polarised during migration, undergoing pulsed contractions in the cell back, while the front protrudes. Contractions then re-localise to the cell centre, displaying radial polarity during constriction. Thus, behavioural change correlates with a change in the polarity of the contractile cytoskeletal network. To explore how manipulating actomyosin contractility affects the contractile network, we interfered with Rho1, Rok and Myosin phosphatase. We show that cellular contractility levels impact on the behaviour of the contractile network, with pulsed contractions only occurring in cells with intermediate contractility levels. Interestingly, increasing Rho1 levels interferes with pulsed contractions, instead leading to a cycling of cells between two states characterised by a junctional cortical and an apicomedial actin network. Thus, increasing contractility is sufficient to impact on cytoskeletal architecture and, consequently, cell behaviour. Additionally, we show that apical constriction can take place without pulsed contractions, raising questions about the roles of pulsed contractions in constriction.

## Results

### LECs undergo periodic apical contractions

To gain insights into the LECs’ switch from migration to constriction, we analysed the behaviour of their actin cytoskeleton. We imaged LECs using *in vivo* 4D microscopy of the F-actin marker GMA-GFP, an actin-binding fragment of moesin fused with GFP (Bloor and Kiehart, 2001). GMA-GFP revealed a dynamic apicomedial actin network that contracted periodically (Figs. 1B; S1A). We observed ‘flows’, where GMA-GFP fluorescence moved through the cell, and ‘foci’, where fluorescence coalesced in distinct regions (Figs. 1B; S1B’; Movie S1). GMA-GFP also labelled the junctional cortical actin at the cell-cell interfaces (Fig. 1B). We used F-actin labelling to explore contractility of the actin network, as it visualises the result of a contractile event (assembly of a focus).

### The activity of the pulsatile network correlates with LEC behaviour

Contractile behaviour correlated with four distinct phases of LEC behaviour (Fig. 1C; Movie S2): *Phase 0* – Stationary LECs without visible cytoskeletal activity. *Phase 1* – During early migration, LECs created a lamellipodium and migrated posteriorly; the cytoskeleton showed diffuse apical activity. *Phase 2* – During late migration, LECs produced a lamellipodium at the front and two actin foci in the back (Fig. 1B-E). The individual actin foci assembled with a period of 180±0.7s (n=7) and contractility alternated between the two foci with around half their individual period (90±2.4s; n=7). *Phase 3* – Constricting LECs showed one central actin focus (Fig. 1B-E) with a pulse period comparable to foci in migrating LECs (180±3.3s; n=7). The pulse period did not increase as constriction progressed (late phase 3; 180±0.4s; n=4).

The transition between phases 2 and 3 was characterised by lamellipodium disappearance and loss of two actin foci, with one focus appearing in the cell centre (Fig. 1C,E). For the duration of the transition, pulsing became more irregular (n=4/7; Fig. 1E) and actin foci and flow patterns changed (n=5/7; Fig. S1B”).

In LECs which alter migration direction from posterior to dorsal (Fig. 1 A), the actin foci reorganised to the new cell back. Eventually, the cells ceased migration and constricted with one central focus (n=3) (Fig. 1F; Movie S3).

Cytoskeletal behaviour was not an artefact of GMA-GFP expression, as the actin marker LifeActin-Ruby (Hatan et al., 2011) allowed similar observations (Figs. 1D; S1C).

Cytoskeletal activity was not only correlated with cell behaviour, but also with cell shape. During migration, cells tended to be elongated along the dorso-ventral (d-v) axis, with actin foci in the back, whereas during constriction cells were rounder, with a central actin focus (Fig. 1G). To become rounder, cells mainly changed shape along the d-v axis (Fig. S1D).

Overall, pulsed contractions of the LEC actin cytoskeleton adopt a highly coordinated pattern, which correlates with cell behaviour.

### Apical area fluctuations go through distinct phases

We next assessed the role of pulsed contractions in LEC behaviour. Pulsed contractions are crucial in driving apical constriction (Levayer and Lecuit, 2012; Martin et al., 2009; Mason and Martin, 2011; Solon et al., 2009). If this were the case in LECs, we would expect the rhythmic activity of actin foci to be correlated with rhythmic changes in apical cell area. We therefore quantified the apical area change in the different phases of LEC behaviour (Fig. 2A).

**Figure 2.**
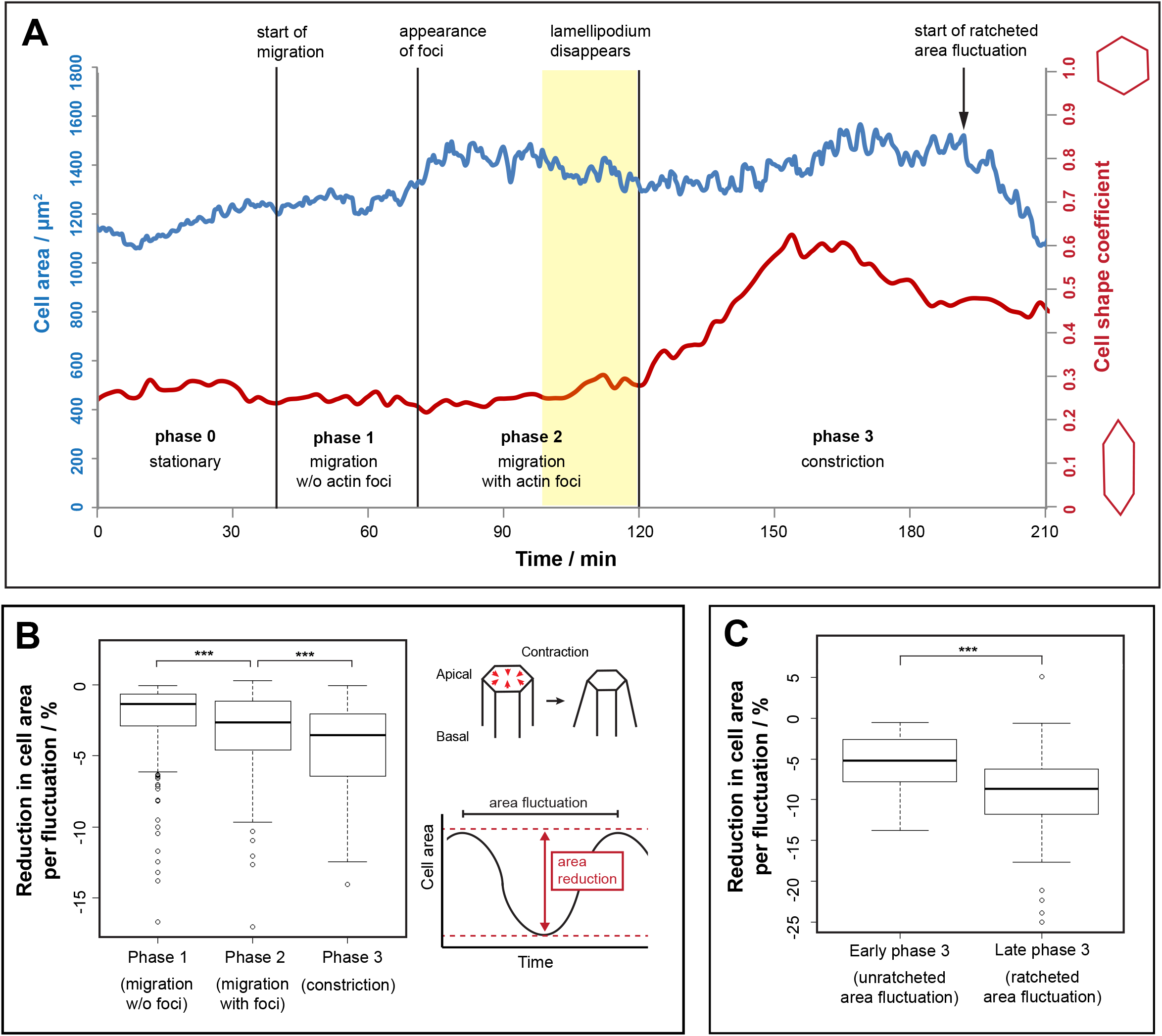
LECs undergo cell area fluctuations that correlate with their behaviour. (**A**) Tracking of LEC area over time. Cell area (blue) and cell shape coefficient (red) shown. Extent of cell area fluctuations increases as cell goes through the four behavioural phases. Cell become rounder when lamellipodium disappears. (**B**) Reduction in cell area per area fluctuation in percent (scheme on right) in phases 1, 2 and 3 (***p<0.001; n=7). (**C**) Reduction in cell area per area fluctuation during unratcheted *vs*. ratcheted constriction (***p<0.001; n=4).

During early migration, no foci were visible, but apical cell area was fluctuating with an average reduction in area by −1.4±0.1% (n=7; Fig. 2B). With the appearance of foci, fluctuations increased significantly (−2.6±0.2%; n=7; Fig. 2B). During constriction, area fluctuations further increased (−3.6±0.3%; n=7; Fig. 2B). Constriction can be divided into an ‘early’ phase, in which the apical area was not reduced but changed its shape, with cells becoming rounder (Figs. 1G; 2A; S1D), and a ‘late’ phase, in which apical area decreased (Fig. 2A). This ‘late’ constriction was characterised by a further significant increase in apical area fluctuation (Fig. 2C) as well as ratcheted constrictions, which ultimately led to delamination. Thus, the ‘early’ constriction phase can be considered a transition phase in which LECs alter their apical shape and undergo pulsed contractions but do not considerably reduce apical area (as they do later on).

This sequence of behavioural changes resembles embryonic cells during gastrulation, which go through comparable phases of unconstricting, unratcheted and ratcheted contractions (Roh-Johnson et al., 2012; Xie and Martin, 2015). In LECs, however, unconstricting area fluctuations take place without detectable pulsatile behaviour, and all observed pulsed contractions lead to area fluctuations.

### Pulsed contractions drive apical area fluctuation

We next asked whether pulsed contractions correlate with cell area fluctuations (Fig. 3A). We found that, in 51% of fluctuations in migrating LECs and in 74% of fluctuations in constricting LECs, one actin focus occurred during one fluctuation (Fig. 3B). We also found that foci appeared around 30±5.3s (n=118 foci) before LEC area was smallest (Fig. 3C). This suggests that area fluctuations correlate with actin foci and that the contractile event drives cell area reduction.

**Figure 3.**
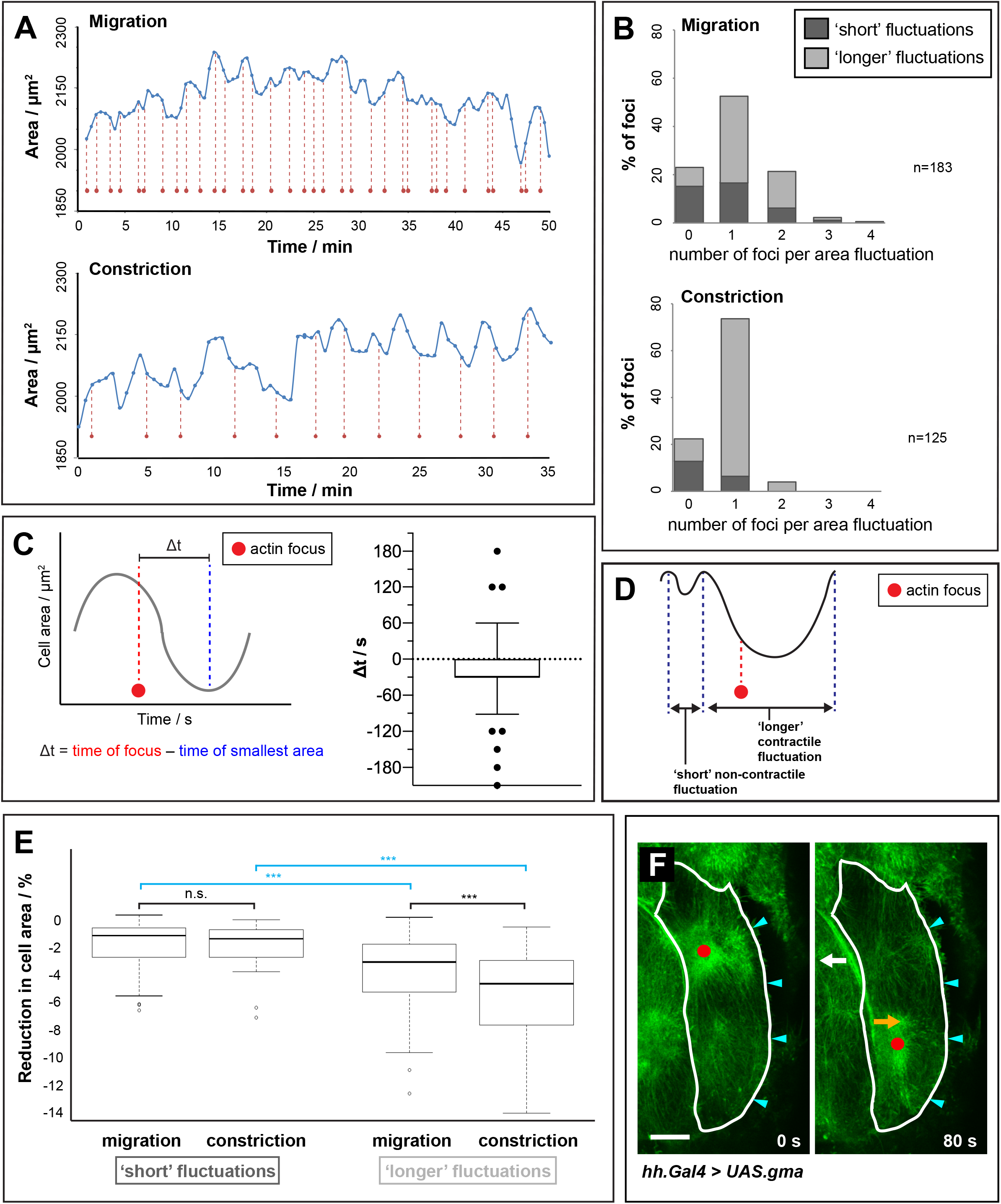
Reduction in LEC area correlates with occurrence of actin foci. (**A**) Cell area fluctuation and actin foci (red dots) shown over time. During constriction, foci appear more regularly, one focus roughly corresponding with one fluctuation. (**B**) Analysis of time difference between occurrence of actin focus and smallest LEC area (Δt). Left: scheme depicting Δt. Right: Δt (Δt=30±5.3s; 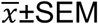; n=188 foci). (**C**) Number of actin foci per area fluctuation. Bars subdivided depending on duration of fluctuation – ‘short’ (≤90s) and ‘longer’ fluctuations (>90s). During migration, LECs show higher variability in foci number per fluctuation. Migrating cells also show higher proportion of ‘short’ fluctuations. Most fluctuations without foci are ‘short’. (**D**) Scheme illustrating ‘short’ non-contractile and ‘longer’ contractile fluctuation. (**E**) Reduction in cell area per pulsed contraction during migration and constriction, for ‘short’ and ‘longer’ fluctuations (***p<0.001; n=7). (**F**) Migrating LECs constrict asymmetrically. Migrating LEC undergoing two consecutive contractions shown. GMA-GFP labels F-actin. Red dot, actin focus; cyan arrowheads, lamellipodium. Cell outline drawn at 0s and the same outline shown at 80s. At 80s, cell expands in region of the focus present at 0s (white arrow), whereas it contracts in region of the new focus (orange arrow). Bar, 10 μm.

Besides area fluctuations that correlated with foci, we also observed fluctuations without foci (Fig. 3B). We hypothesised that these fluctuations might be due to external forces exerted by contracting neighbouring cells, which could interfere with regular fluctuations. Such events might have a shorter duration and lead to smaller area reduction than pulsed contractions (Fig. 3D). Categorising individual fluctuations by duration, we found that the majority of fluctuations not coinciding with an focus were ‘short’ (≤90s) (Fig. 3B). These ‘short’ fluctuations did not reduce the cell area much – the reduction, both during migration (−1.19±0.2%; n=7) and constriction (−1.41±0.4%; n=7) (Fig. 3E), was comparable to that in LECs that migrated without visible pulsatile activity (−1.36±0.1%; p=0.64). However, for ‘longer’ fluctuations (>90s), there was a significant difference in area reduction per fluctuation between migration and constriction (Fig. 3E). Overall, this suggests that the majority of area fluctuations that occur without an accompanying actin focus are ‘short’ non-contractile fluctuations that might be due to external forces by neighbouring LECs.

Furthermore, in migrating LECs, the correlation between area fluctuations and actin foci was less strong than in constricting LECs – around 25% of the fluctuations in migrating LECs showed two foci, and overall the number of ‘short’ fluctuations involving foci was higher than in constricting LECs (Fig. 3B). The weaker correlation could be due to the two alternating contractile events in different cell regions affecting cell shape change unevenly (Fig. 3F).

Taken together, our observations suggest that apicomedial network contraction reduces LEC area. Both Sqh::GFP (Royou et al., 2004) and Rok::GFP (Abreu-Blanco et al., 2014) co-localised with foci labelled with LifeActin-Ruby (Fig. 4A,B). This corroborates the notion that network contractility is created by actomyosin activity.

**Figure 4.**
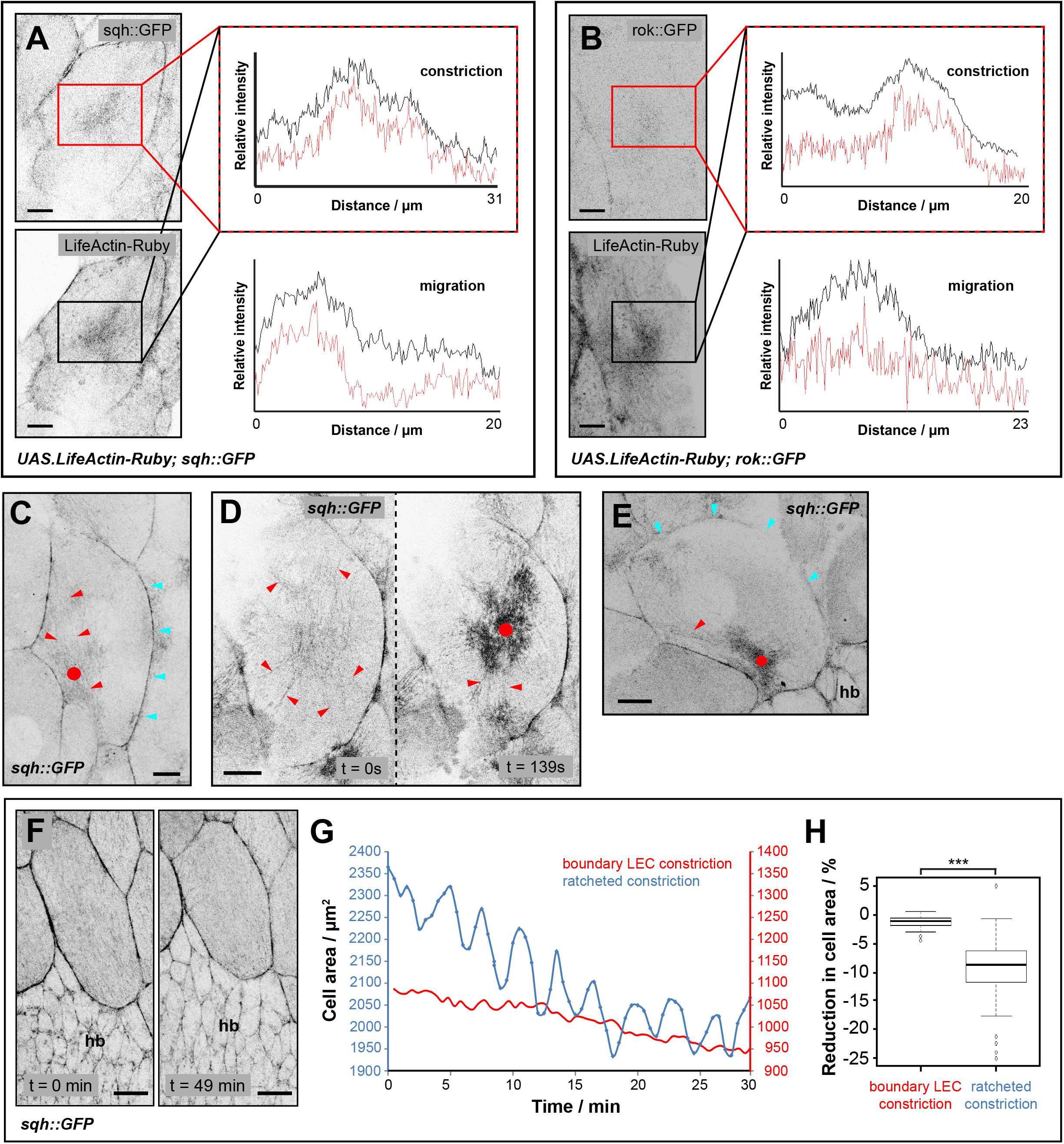
Dynamic behaviour of the LEC cytoskeleton. (**A,B**) LifeActin-Ruby co-localises with (**A**) Sqh::GFP and (**B**) Rok::GFP in actin foci and cell-cell interfaces, during migration and constriction. Plot profiles of relative fluorescence intensity in rectangular region shown. (**C-E**) Sqh::GFP labels actin foci (dots), actin bundles (red arrowheads) and cell-cell interfaces. Cyan arrowheads, lamellipodium. (**C**) Posteriorly migrating LEC, showing actin focus and bundles in back as well as lamellipodium in front. Bundles are organised along d-v axis. (**D**) Constricting LEC before (left) and during (right) pulsed contraction. Actin bundles are organised radially. (**E**) Dorsally migrating LEC, showing actin focus and bundles in back and lamellipodium at front. (**F-H**) Constriction of boundary LECs during early morphogenesis. (**F**) Sqh::GFP labels cell-cell interfaces. Also some diffuse labelling all over apical cell area, but no actin foci visible. LEC constricts over time. (**G**) Comparison of area fluctuations over time between LEC undergoing ratcheted constriction and constricting boundary LEC, which shows little area fluctuation. Note that boundary cell constricts rather slowly. (**H**) Reduction in cell area per pulsed contraction during ratcheted constriction and early boundary LEC constriction (***p<0.001; n=4). All bars, 10 μm; hb, histoblasts. Anterior, left; dorsal, up.

### LECs show distinct cytoskeletal architecture during migration and constriction

Sqh::GFP, Rok::GFP, LifeActin-Ruby and GMA-GFP not only labelled foci, but also localised to the cell cortex at cell-cell junctional interfaces (Figs. 1B; 4A,B). In early stationary and migratory LECs, Sqh::GFP levels tended to be higher at anterior-posterior (a-p) interfaces, compared to d-v interfaces (Fig. S2A’). Constricting cells, however, showed no differences in Sqh::GFP localisation between a-p and d-v interfaces (Fig. S2A”).

In addition, Sqh::GFP and GMA-GFP also labelled actin bundles. Migrating LECs displayed actin bundles in their back, where also myosin/actin foci occurred (Figs. 4C; S2B,C). These bundles resembled stress fibres in migrating fibroblasts (Pellegrin and Mellor, 2007). Deformation of cells where actin bundles meet the cell-cell junctions suggests that the bundles can exert pulling forces (Fig. S2D). In constricting LECs, actin bundles appeared to connect a contractile apicomedial network radially to the cell-cell junctions (Fig. 4D; Movie S4). After repolarisation, actin bundles, like foci, were found in the new back of the now dorsally migrating cells (Fig. 4E). Thus, besides foci localisation, other aspects of cytoskeletal architecture also correlate with cell behaviour and polarity, which changes as cells transit from migration to constriction.

### Not all LECs undergo pulsed contractions

Studying Sqh::GFP dynamics, we noticed that those LECs which border histoblast nests at the beginning of morphogenesis (boundary LECs (Teng et al., 2017)) constrict apically without displaying actin foci. Instead, they show Sqh::GFP at their cell-cell junctions, as well as diffuse labelling across their apical surface (Fig. 4F; Movie S5). When constricting, boundary LECs showed little apical area fluctuations, compared to LECs undergoing pulsed contractions (Fig. 4G,H). This highlights that not all LECs that reduce their apical area do so while contracting in a pulsatile manner.

### Reducing actomyosin contractility

To gain insights into the regulation of pulsed contractions, we interfered genetically with Rok and the Myosin-binding subunit (Mbs) of Myosin phosphatase. From previous studies, we expected both RNAi knockdown of Rok and constitutive activation of Mbs to reduce the amount of phosphorylated Myosin II and thus cell contractility (Dawes-Hoang, 2005; Fischer et al., 2014; Lee and Treisman, 2004; Munjal et al., 2015; Valencia-Expósito et al., 2016; Vasquez et al., 2014). In LECs, overexpression of a constitutively active form of Mbs (MbsN300 (Lee and Treisman, 2004)) impairs apical constriction (Ninov et al., 2007).

Rok-RNAi did not interfere with LEC migration (Fig. S3A,B), but LECs showed a phenotype indicative of reduced contractility. Cells had an increased apical area compared to controls (Fig. 5A,B) and apical area fluctuations were significantly reduced (Fig. 5C). In the back of migrating cells, actin accumulated periodically, travelling along the membrane in d-v direction (Fig. 5A; Movies S6, S7). These contractile flows only occurred basolaterally underneath protruding lamellipodia of neighbouring cells. Thus, these contractions might be a reaction to pushing forces created by the lamellipodia of neighbours. Occasionally, we observed this behaviour also in wild-type LECs (Fig. 5D). Rok-RNAi might increase this phenomenon, as lamellipodia push more strongly on top of their neighbours due to their reduced contractility.

**Figure 5.**
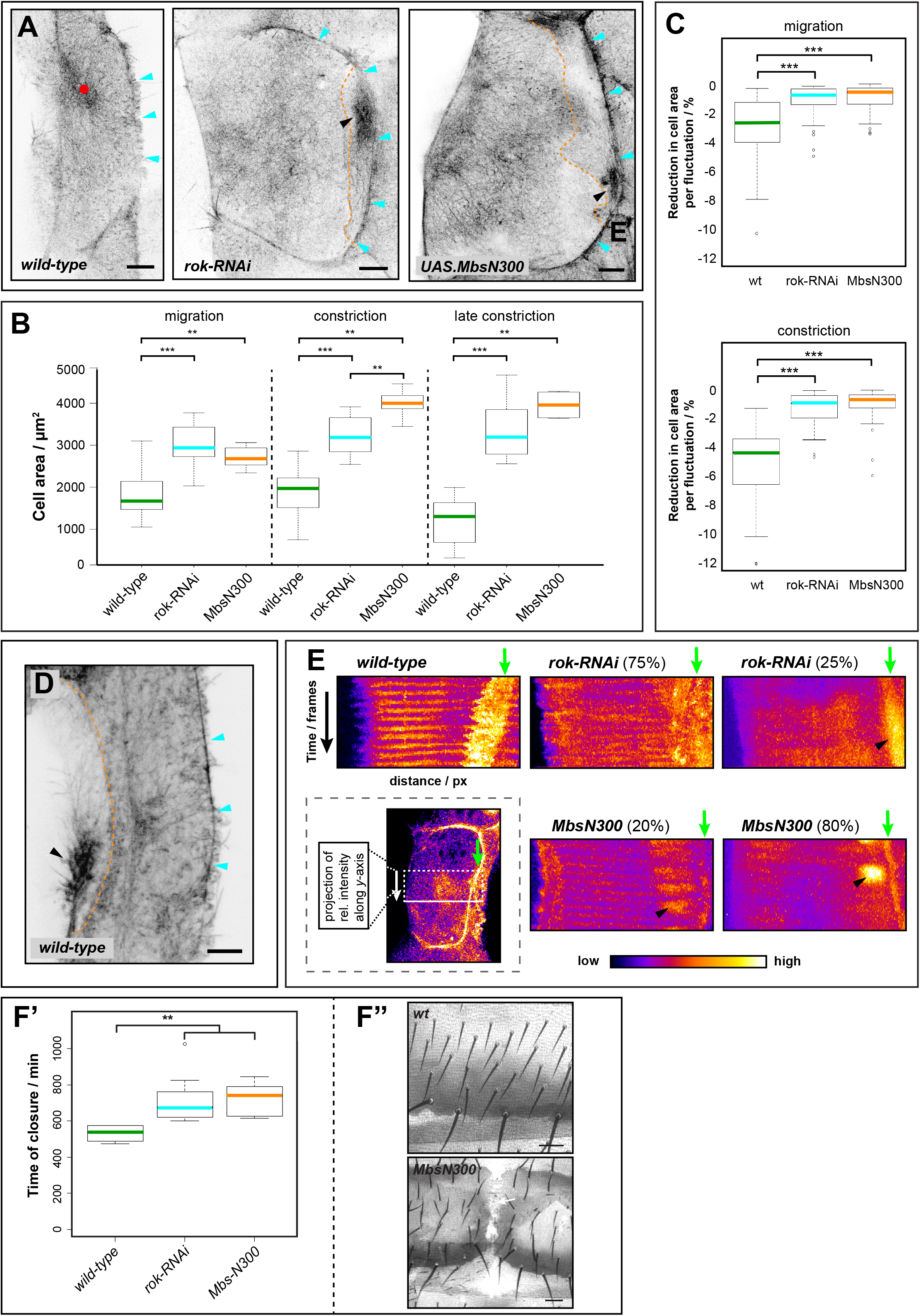
Reduction in LEC contractility interferes with actin foci formation, cell shape and area fluctuation. (**A**) Wild-type, *rok-RNAi* and *UAS.MbsN300* LECs during migration. GMA-GFP labels F-actin. Cells generate lamellipodium (cyan arrowheads). Wild-type LEC shows actin focus (dot), whereas *rok-RNAi* and *UAS.MbsN300* LECs show more diffuse cytoskeleton labelling without foci. Neighbours create contractile flows in their back (black arrowheads, dotted line outlines overlap between cells). Bars, 10 μm. (**B**) Apical area of *wild-type, rok-RNAi* and *MbsN300* LECs during migration, early constriction and late constriction. (**C**) Reduction in cell area per pulsed contraction in wild-type, *rok-RNAi* and *MbsN300*. (**D**) Migrating wild-type LEC creates contractile flow in its back (black arrowhead). GMA-GFP labels F-actin. Bar, 10 μm. (**E**) Kymograph of LEC cell centre during constriction. Kymograph shows *y*-projection of rectangular region of interest (see scheme in box) over time. GMA-GFP labels F-actin. Wild-type LEC shows rhythmic actin activity. Due to area fluctuation, membrane region appears broad (green arrow). In 75% of *rok-RNAi* and 20% of *MbsN300* LECs, some rhythmical activity visible, but more diffuse than wild-type. In 25% of *rok-RNAi* and 80% of *MbsN300* LECs, no rhythmical activity visible. Contractile flows in back of neighbouring cell visible (black arrowhead). In mutants, membrane region appears thin due to fewer area fluctuations (green arrows). (**F**) *rok-RNAi* and *MbsN300* prolong time of abdominal closure leading occasionally to dorsal cleft phenotype. (**F’**) Timing of abdominal closure. Wild-type, *rok-RNAi* and *MbsN300* shown. (**F”**) Wild-type cuticle and dorsal cleft phenotype in MbsN300 pupa. Bars, 50 μm. *** p<0.001; ** p<0.01.

During constriction, Rok-RNAi had an effect on LEC contractility. LEC apical area was increased significantly (Fig. 5B), and area fluctuations were decreased (Fig. 5C). Fluctuations (−0.59±0.1% cell area; n=8) were even smaller than in early migrating wild-type LECs, which do not show pulsed contractions (−1.36±0.1%; n=7; p<0.001).

In addition, Rok-RNAi affected pulsed contractions. 75% of analysed LECs showed some actin foci, whereas in 25%, foci were absent (n=8; Fig. 5E). Where foci could be observed, their pulsation period was comparable to wild-type (180s±1.7s; n=5; p=0.11), but foci were more diffuse (Fig. 5E). In strong phenotypes, GMA-GFP labelled a less dynamic apicomedial network, which did not generate any foci and showed some diffuse activity (Fig. 5A,E; Movie S7).

Overexpression of MbsN300 showed a similar phenotype to Rok-RNAi (Figs. 5A-C,E,F; S3B; Movie S8). However, the MbsN300 phenotype was stronger, with a higher proportion of ‘strong’ phenotypes, which did not generate any actin foci (Fig. 5E).

To ask whether defects in LEC contractility impact on morphogenesis, we assessed the time of abdominal closure in control, Rok-RNAi and MbsN300 pupae. We found that in both experiments, closure was delayed (Fig. 5F’). In some cases, this led to closure defects, where LECs did not complete morphogenesis successfully (2.6% of Rok-RNAi pupae (n=39); 60.8% of MbsN300 pupae (n=51); Fig. 5F”). This suggests that impaired contractility can lead to morphogenesis defects, but that in many cases, LECs still delaminate.

Our data show that activity of both kinase and phosphatase (which activate and deactivate myosin II, respectively) is crucial for cell contractility, pulsed contractions and apical area fluctuations. However, even without detectable actin foci and thus significantly less apical area fluctuation, LECs can constrict and delaminate successfully.

### Increasing actomyosin contractility

Next, we asked what effect an increase in contractility has on LEC pulsatile activity. In the *Drosophila* embryo, constitutive activation of Sqh or Rho1 interferes with the activity of the pulsatile actomyosin network (Fischer et al., 2014; Mason et al., 2016; Munjal et al., 2015; Valencia-Expósito et al., 2016; Vasquez et al., 2014).

Firstly, we overexpressed in LECs a constitutively active form of Rok (Rok-CAT), which leads to excessive phosphorylation and thus activation of myosin II (Verdier et al., 2006; Warner and Longmore, 2009). Using Gal80ts to repress Rok-CAT expression before pupal stages allowed us to assess different levels of Rok-CAT expression by inactivating the repressor either efficiently at 29°C for strong expression or inefficiently at 25°C for weaker expression. This approach resulted in two distinct phenotypes: (1) In ‘weak’ phenotypes, LECs showed migratory and constrictive behaviour (Fig. 6A,B; Movie S9; 100% of 25°C experiments, n=5; 35.3% of 29°C experiments, n=17). Like in wild-type, migrating LECs displayed periodic actin foci in their back, which moved to the centre during constriction (Fig. 6B) and led to area fluctuations (Fig. 6C). Noticeably, LECs underwent delamination, which was characterised by a more pronounced contractile ring compared to wild-type (Fig. 6A; Movie S9). Furthermore, constricting cells showed blebbing at the apical junctional cortex (Fig. 6A; Movie S9). (2) In ‘strong’ phenotypes, LECs did not migrate and only constricted (Fig. 6D,E; Movie S10; 64.7% of 29°C experiments, n=17). Constricting cells showed extensive blebbing and formed cortical actin bundles apically, as well as stress-fibre-like bundles more basally (Fig. 6D). In addition, cells did not show pulsatile activity (Fig. 6E) and lacked notable area fluctuation (Fig. 6C).

**Figure 6.**
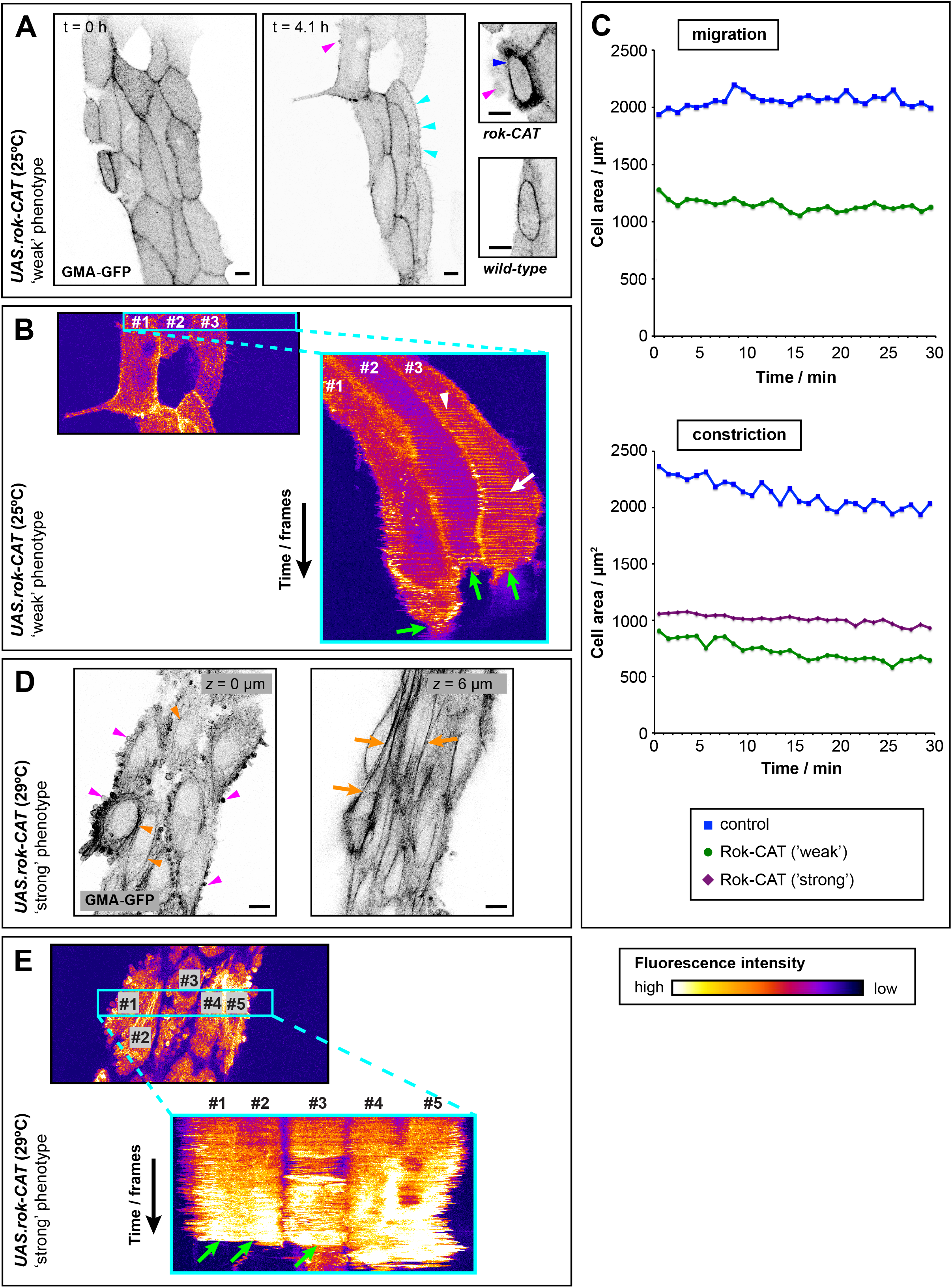
Increasing LEC contractility by expression of constitutively active Rok (Rok-CAT). GMA-GFP labels F-actin. (**A,B**) ‘Weak’ phenotype; transgene expression at 25°C. (**A**) Start of morphogenesis (left) and LEC migration (right). LECs migrate, showing some blebbing (magenta arrowheads) and lamellipodia (cyan arrowheads). Details: Compared to wild-type, constricting Rok-CAT cell shows high cortical actin (blue arrowhead) and blebbing. (**B**) Kymograph of three LECs (right panel). Projected region indicated in left panel. Cells move to the right, eventually cease to move, constrict and delaminate (green arrows). In cell #3, contractile activity is in back during migration (white arrowhead) and in centre during constriction (white arrow). (**C**) Comparing cell area over time for wild-type, ‘weak’ and ‘strong’ Rok-CAT phenotype. Part of migration and constriction phase shown. Wild-type and ‘weak’ Rok-CAT LECs show cell area fluctuations, but ‘strong’ Rok-CAT LEC does not. Representative cell shown for each experiment. (**D,E**) ‘Strong’ phenotype; transgene expression at 29°C. (**D**) Constricting LECs. Left: Apically, LECs show extensive blebbing (magenta arrowheads) and cortical actin bundles (orange arrowheads). Right: More basally, cells generate extensive stress fibre-like actin bundles (orange arrows). (**E**) Kymograph of five LECs (bottom panel). Projected region indicated in top panel. No rhythmical cytoskeletal activity. Cells eventually delaminate (green arrows). All bars, 10 μm.

Secondly, we overexpressed in LECs a constitutively active form of Rho1 (Rho1-CA (Fanto et al., 2000)). We have shown previously that Rho1-CA overexpression leads to loss of migration and defects in abdominal morphogenesis (Bischoff, 2012). We hypothesised that overexpressing Rho1-CA should lead to more Rok being activated and thus a phenotype similar to Rok-CAT. Indeed, Rho1-CA phenotypes resembled Rok-CAT phenotypes, with cells displaying cortical blebbing, junctional cortical actin localisation and absence of rhythmic cytoskeletal activity (Fig. 7A,B; Movies S11,S12), which is reflected in the lack of cell area fluctuation (Fig. 7C). However, unlike Rok-CAT cells, Rho-CA LECs did not create excessive cortical actin bundles (Fig. 7A,B). With respect to cell migration, Rho1-CA pupae showed two phenotypes: ‘migrating’ phenotypes (29%, n=7; Fig. 7A; Movie S11), and ‘non-migrating’ phenotypes (71%), in which LECs only constricted (Fig. 7B; Movie S12). Unlike Rok-CAT cells, none of the analysed Rho-CA cells showed pulsed contractions (n=7).

**Figure 7.**
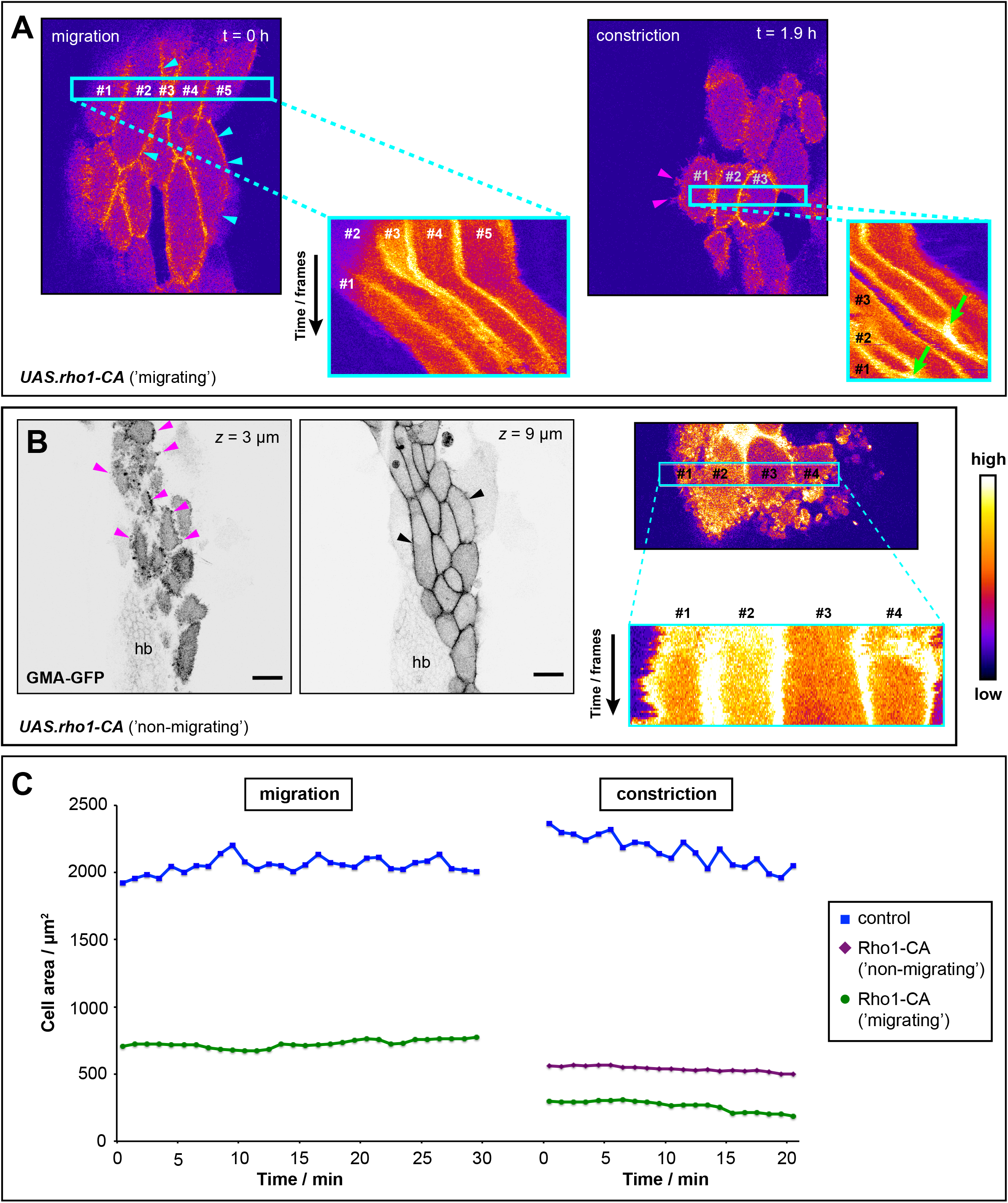
Increasing LEC contractility by constitutive activation of Rho1 (Rho1-CA). (**A,B**) GMA-GFP labels F-actin. (**A**) ‘Migrating’ phenotype. Kymograph of LECs during migration and constriction. No rhythmical cytoskeletal activity visible. Projected region indicated in overview images. Cells migrate (lamellipodia, cyan arrowheads), eventually constrict showing cortical blebbing (magenta arrowheads) and delaminate (green arrowheads). (**B**) ‘Non-migrating’ phenotype. Left panels: Micrographs of apical and lateral *z*-section. Constricting cells show extensive apical cortical blebbing (magenta arrowheads) and actin labelling at cell interfaces (black arrowheads. Bars, 20 μm. Right panels: Kymograph of four LECs. Projected region indicated in overview image (top panel). No rhythmical cytoskeletal activity visible. (**C**) Comparing cell area over time for wild-type, ‘migrating’ and ‘non-migrating’ Rho1-CA LEC. Part of migration and constriction phase shown. Wild-type LEC shows area fluctuations, but both ‘migrating’ and ‘non-migrating’ Rok-CAT LECs do not. Representative cell shown for each experiment.

These data show that increasing contractility can lead to the loss of migratory behaviour and interfere with pulsed contractions and apical area fluctuations. Expressing Rho-CA in all LECs results in abdominal cleft phenotypes in 63% of cases (Bischoff, 2012). Thus, although apical constriction is impaired, LECs delaminate successfully in many cases.

### Increasing levels of Rho1 induces rhythmical remodelling of the apicomedial network

Besides activating contractility by expressing constitutively active forms of Rok and Rho1, we also increased the amount of wild-type Rho1 in LECs (*UAS.rho1* (Prokopenko et al., 1999)). Interestingly, these cells did not undergo pulsed contractions, but cycled between two distinct states characterised by (1) the presence of apicomedial actin (Fig. 8A,B) and (2) the absence of apicomedial actin but increased junctional cortical actin and blebbing (Fig. 8A’,B; Movie S13). There was no difference in the mean duration of each state, however, there was considerable variation (16±13.6 min (apicomedial) and 21±35.6 min (junctional cortical); p=0.9797; Fig. 8C). Cells switched between the two states in 2.0±1.0 min (apicomedial to junctional cortical) and 2.6±1.6 min (junctional cortical to apicomedial), respectively. Interestingly, cells delaminated in either state, however, most cells delaminated in the junctional cortical state (Fig. 8D). Depending on the strength of the phenotype, cells were ‘migrating’ (n=3 pupae; Fig. 8B) or ‘non-migrating’ (n=2 pupae; Fig. 8A).

**Figure 8.**
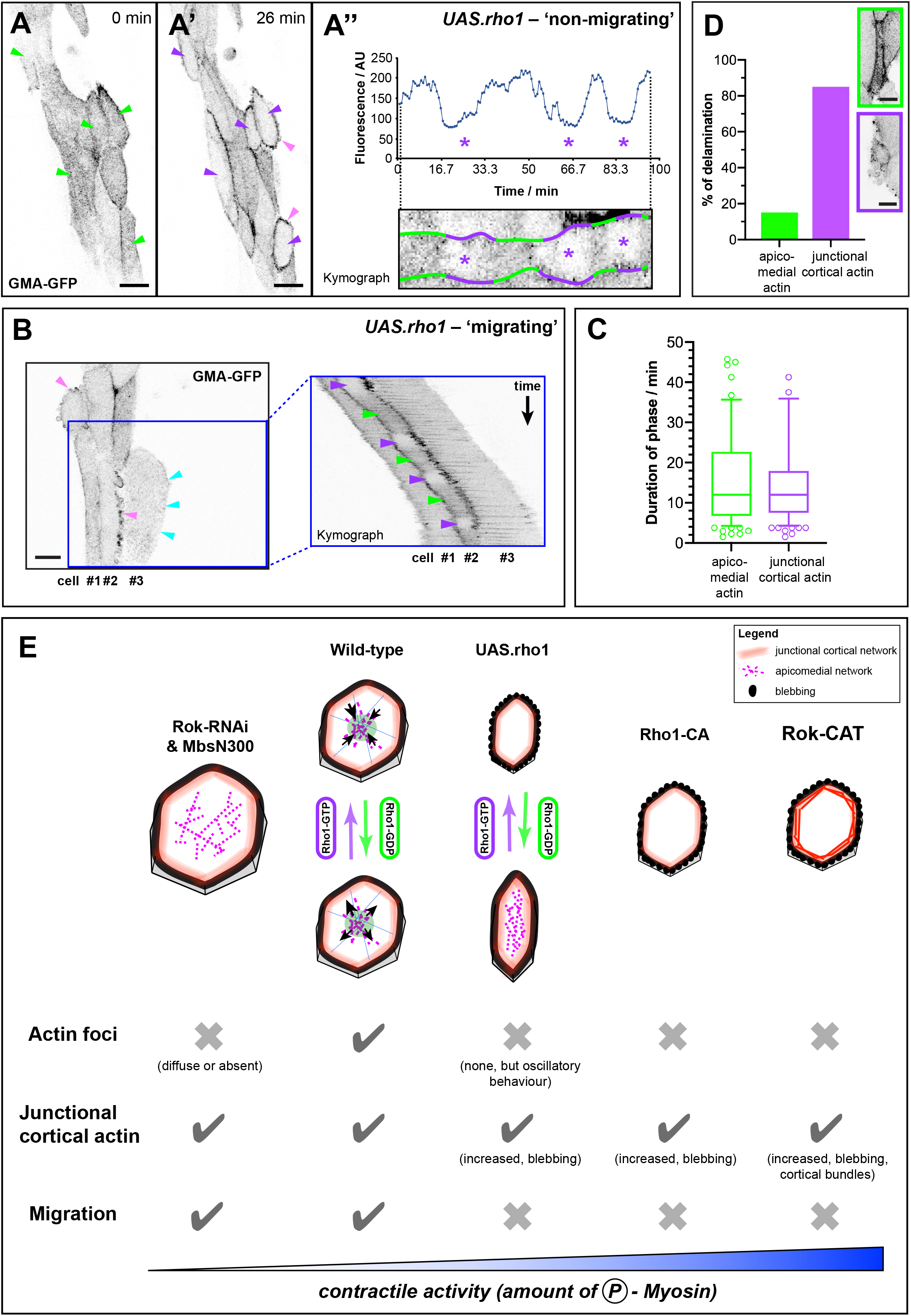
Overexpression of Rho1 leads to oscillatory changes in the cytoskeletal network. (**A,B**) GMA-GFP labels F-actin. Bars, 20 μm. (**A-A”**) ‘Non-migrating’ phenotype. LECs cycle between two states: (**A**) presence of apicomedial actin (green arrowheads) and (A’) absence of apicomedial actin but increased junctional cortical actin (purple arrowheads) and cortical blebbing (pink arrowheads). (A”) Fluorescence intensity of cell centre of cycling LEC (top) and kymograph of same cell (bottom). Purple asterisks indicate junctional cortical states. (**B**) ‘Migrating’ phenotype. Left: LECs forming lamellipodia (cyan arrowheads) and blebs (pink arrowheads). Right: Kymograph from area indicated by blue box. Cells migrate posteriorly. Cell #2 cycles between apicomedial (green arrowheads) and junctional cortical actin state (purple arrowheads). (**C**) Duration of the two states. Eight outliers excluded in plot. (**D**) LECs delaminate in both states. Top box, cell delaminating with apicomedial actin and no blebbing; note that actin is increased. Bottom box, cell delaminating with junctional cortical actin and blebbing. Bars, 10 μm. (**E**) Cellular contractility coordinates LEC behaviour and cytoskeletal dynamics. Amount of LEC contractility affects presence of pulsed contractions (actin foci), amount of junctional actin, as well as presence or absence of cortical blebbing, and migration. Only with intermediate contractility levels, cells show pulsed contractions. LECs with increased Rho1 levels (*UAS.rho1*) cycle between two distinct states, which do not show pulsed contractions. Reducing contractility (Rok-RNAi, MbsN300) interferes with pulsations and leads to larger cell area. Increasing contractility (Rho-CA) leads to cortical blebbing and interferes with pulsations and migration; F-actin localises to junctional cortex. Very strong increase of contractility (Rok-CAT) additionally leads to formation of cortical actin bundles.

Our results suggest that adding additional Rho1, which can be switched on and off by the cells’ endogenous machinery, causes LECs to cycle between two states characterised by distinct cytoskeletal networks. Thus, rhythmical activity appears not to be limited to pulsed contractions of an actomyosin network, but also extend to more general rhythmical remodelling of cytoskeletal architecture (Fig. 8E).

## Discussion

We show that the apicomedial actin cytoskeleton of LECs undergoes pulsed contractions, while cells transit from migration to constriction (Fig. 1A,B). During this transition, cytoskeletal activity is highly coordinated and correlates with cell behaviour and shape (Fig. 1B-G). Pulsed contractions are correlated with cell area fluctuations (Figs. 2; 3) and LECs go through distinct phases of contractile activity (Fig. 1C; 2A).

We find that a cell’s level of contractile activity determines the behaviour of its actin cytoskeleton, as well as cytoskeletal architecture and cell behaviour (Fig. 8E). Not only are pulsed contractions dependent on intermediate levels of contractility, but moderately increasing contractility causes LECs to cycle between two states, characterised by a junctional cortical and an apicomedial actin network, respectively (Fig. 8). Moreover, our data suggest that constriction can occur without pulsed contractions, raising questions about their contribution to constriction.

### The LEC cytoskeleton is highly organised and undergoes dynamic changes during behavioural change

Our study highlights the complexity of a cell’s cytoskeleton during morphogenesis. Migrating LECs are planar polarised, with their apical actin cytoskeleton protruding at the front and undergoing pulsed contractions in the back (Fig. 1B-E). Furthermore, LECs create actin bundles in their back (Figs. 4C; S2B,C) and Sqh::GFP preferentially localises to a-p cell-cell junctions (Fig. S2A’), similar to embryonic cells during germband extension (Zallen and Wieschaus, 2004). As LECs constrict, their cytoskeleton becomes radially polarised, with a central actin focus (Figs. 1B-E; 4D). They also show radial actin bundles that appear to connect the apical network to cell-cell junctions, resembling a spider’s web (Fig. 4D). In addition, Sqh::GFP is distributed evenly at all interfaces (Fig. S2A”). Thus, a crucial step during LEC behavioural change is a change in cell polarity from PCP to RCP that underlies a change in cytoskeletal activity as well as an overall reorganisation of cytoskeletal architecture (Fig. S4). Pulsed contractions are not involved in this change, as Rho-CA cells, which do not pulse, also transit from migration to constriction (Movie S11).

What regulates this polarity change requires further investigation. We have previously shown that LEC polarity depends on PCP signalling (Bischoff, 2012). Cytoskeletal asymmetry in migrating LECs could be due to localised activity of small Rho GTPases, with Rho1 controlling pulsed contractions in the cell back and Rac1 promoting lamellipodia formation at the front. Such mutually exclusive localisation has been shown in various cell types (Cao et al., 2015; Ohta et al., 2006; Sander et al., 1999). In addition, several studies have found asymmetric localisation of Rho1 activity (Simoes et al., 2006; van Impel et al., 2009) and reported Rho1 downstream of PCP signalling (Carmona-Fontaine et al., 2008; Tree et al., 2002). We show that Rok::GFP co-localises with actin foci (Fig. 4B) and that manipulating both Rok and Rho1 activity interferes with LEC migration and pulsed contractions (Figs. 5–8), suggesting that localising Rho1 activity is crucial for coordinating behavioural change.

### Generation of pulsatile activity during behavioural change

Unlike other processes in which cells show pulsed contractions, in LECs actin dynamics involves the formation of distinct areas of contractile activity, depending on LEC behaviour and polarity. Constricting LECs create a single contractile event in the cell centre, while migrating LECs generate two alternating foci in their back (Fig. 1B-E). This difference cannot easily be explained by a change in cellular polarity. Instead, it could be due to different shapes of the apical cell area in migrating and constricting cells. In migrating cells, which are elongated along the d-v axis (Figs. 1G; S1D), one contractile event might not suffice to constrict the whole apicomedial network (Fig. 3F). Constricting cells, however, are rounder (Figs. 1G; S1D), which might allow radial actin recruitment over the whole apical area (Fig. 4D). Alternatively, the two foci could be a consequence of the cytoskeletal architecture of a migrating cell, as seen for myosin localisation in keratinocytes (Wilson et al., 2010). The actin bundles in the back (Figs. 4C; S2B,C) might promote the formation of two lateral foci by inhibiting foci more centrally. Pulsed contractions are not part of the migratory machinery, as early migrating LECs do not show actin foci (Movie S2) and Rho-CA LECs migrate without pulsatile activity (Fig. 6A,B). Interestingly, LECs that create actin foci migrate faster than LECs of an earlier stage, which do not undergo pulsed contractions (Fig. S5A).

Overall, our results suggest that cytoskeletal architecture and cell area determine the localisation and behaviour of contractile events. In addition, comparing pulsatile behaviour in migrating and constricting LECs corroborates the notion that autonomous network properties regulate foci (Banerjee et al., 2017; Munjal et al., 2015), rather than external input from a pulse generator. The two foci in migrating LECs alternate around every 90s. Interestingly, their individual pulse period is around 180s, i.e. similar to the period of central foci during constriction. If there were an external pulse generator, its frequency would need to halve, which appears less plausible than an autonomous assembly of foci.

### The level of contractility affects cytoskeletal dynamics

Altering levels of contractility in LECs showed that pulsed contractions depend on Rok, Myosin phosphatase and Rho1 activity. In this respect, LECs resemble other cells that undergo pulsed contractions (Fischer et al., 2014; Mason et al., 2016; Munjal et al., 2015; Valencia-Expósito et al., 2016; Vasquez et al., 2014). As shown for *Drosophila* germband cells (Munjal et al., 2015) and amnioserosa cells (Fischer et al., 2014), our results indicate that cytoskeletal network dynamics depends on the level of cell contractility (Fig. 8E). Only with intermediate, wild-type levels of contractility did LECs show pulsed contractions. Reducing contractility (Rok-RNAi or MbsN300) interfered with pulsations (Fig. 5A,E), as did increasing contractility (Rho1-CA or Rok-CAT) (Figs. 6E, 7A,B). As in amnioserosa cells (Fischer et al., 2014), reducing LEC contractility also increased cell area (Fig. 5A,B), suggesting that certain contractility levels are needed to maintain normal cell shape.

### The level of contractility affects cytoskeletal architecture and cell behaviour

Besides interfering with pulsed contractions, increasing contractility in LECs had a more far-reaching impact on cytoskeletal architecture. (1) Overexpression of Rho1-CA resulted in F-actin disappearing apicomedially and localising mostly to the junctional cortex (Fig. 7A,B). A similar phenotype has been observed during *Drosophila* gastrulation (Mason et al., 2016), suggesting that Rho1 is involved in determining the ratio of apicomedial *vs*. cortical contractility (Munjal et al., 2015). (2) While overexpression of Rho1-CA can only activate endogenous Rok, overexpression of Rok-CAT can supply large amounts of activated Rok and thus lead to a stronger and more specific activation of myosin II. In LECs, this not only resulted in F-actin disappearing apicomedially and localising mostly to the junctional cortex, but also in the formation of cortical actin bundles (Fig. 6D). In cell culture, activation of myosin has been shown to create actin bundles during stress fibre formation (Chrzanowska-Wodnicka and Burridge, 1996). (3) Increased junctional cortical contractility in both Rho1-CA and Rok-CAT LECs furthermore induced cell blebbing (Figs. 6D; 7B). Blebbing is driven by strong cortical myosin activation (Paluch et al., 2006). During *Drosophila* dorsal closure, activation of Myosin light chain kinase and Mbs leads to cell blebbing (Fischer et al., 2014).

Altering levels of contractility in LECs also affects cell behaviour directly (Fig. 8E). In ‘strong’ phenotypes of both Rok-CAT and Rho-CA pupae, LECs do not generate lamellipodia-like protrusions and do not migrate (Figs. 6E; 7B). Thus, high levels of activated myosin appear to override migratory behaviour and force cells to constrict.

### Rho1 overexpression induces rhythmical cytoskeletal behaviour distinct from pulsed contractions

Increasing wild-type Rho1 levels proved particularly informative (Fig. 8A-D). Firstly, *UAS.rho1* cells show a phenotype consistent with an increase in contractility, i.e. loss of pulsed contractions (Fig. 8A”) and blebbing (Fig. 8A’,B). This suggests excess endogenous Rho1-activating machinery, which constitutively activates some of the additional Rho1. Secondly, when adding additional Rho1, its rhythmical activation and de-activation via the endogenous machinery still appears to occur, leading to the cycling between a state of higher contractility, where actin accumulates at the junctional cortex and cells show blebbing (Fig. 8A’,B), and a state of lower contractility, where the apicomedial network is present but non-pulsatile (Fig. 8A,B). There are two possible explanations for this phenotype: (1) With increasing levels of activated Rho1, LECs shift their ratio of apicomedial *vs*. junctional cortical actin (and thus contractility) towards cortical. This is reversed, when levels of activated Rho1 decrease. (2) High levels of activated Rho1 (and thus contractility) lead to the rupture of the apicomedial network and only the junctional cortical network remains. When Rho1 levels decrease, the apicomedial network forms again. Our results support the first hypothesis, as we observed not only the disappearance of apicomedial actin, but also the beginning of junctional cortical blebbing. This suggests an increase in contractility in the junctional cortical network, rather than just a collapse of the apicomedial network.

Cells transited between the two states in 2.0±1.0 min (apicomedial to junctional cortical) and 2.6±1.6 min (junctional cortical to apicomedial), respectively. This further argues against a sudden rupture of the apicomedial network. It also shows that switching from one state to the other occurs in the same time frame as a pulsed contraction of the apicomedial network in wild-type LECs (3.0±0.002 min).

Compared to pulsed contractions, the duration of the two states in Rho1 overexpressing LECs is far longer and more irregular (Fig. 8C). This could be due to the endogenous machinery having to deal with large amounts of Rho1 that need to be switched on/off, which unbalances the system.

### The role of pulsed contractions in apical constriction

Although pulsed contractions are affected by the manipulation of contractility, we found that most LECs in Rho-RNAi, MbsN300, *UAS.Rho1*, Rho1-CA or Rok-CAT pupae constricted successfully. In all these cases, pulsed contractility of the apicomedial actin network was impaired, but cells still had actin at their junctional cortex (Figs. 5A; 6D; 7B). This indicates that contractility at the junctional cortex alone can drive apical constriction. Similar observations have been made in amnioserosa cells (Saravanan et al., 2013) and during neural tube formation (Christodoulou and Skourides, 2015). Our observation that boundary LECs at the beginning of morphogenesis constrict without pulsed contractions and without noticeable cell area fluctuation (Fig. 4G) further supports the notion that contractility needed for constriction can be created by the junctional cortical network. Another possibility is that the apicomedial network creates non-pulsatile tension that helps drive constriction. For instance, in boundary LECs, diffuse apical Sqh activity can be observed, which might contribute to apical area reduction (Movie S5).

This raises important questions about the role of pulsed contractions in constriction, as also raised by others (Blanchard et al., 2010; Xie and Martin, 2015). LECs begin pulsed contractions while still migrating (Fig. 1C), at a time when LEC shape changes, and thus tissue remodelling, intensify. Also, for most of morphogenesis, LECs undergo pulsed contractions without constricting, merely changing shape; only when ratcheted constrictions begin, cell area reduces notably (Fig. 2A). This suggests that pulsed contractions do not drive apical constriction *per se*. Instead, they might have other roles, such as helping to maintain cell shape in an environment where the activity of neighbouring cells creates pushing and pulling forces. Ultimately, this will help to maintain tissue integrity during morphogenesis (Coravos et al., 2017). Alternatively, pulsed contractions could cooperate with junctional cortical contractility to create sufficient forces to drive apical constriction more effectively. LEC constriction that is accompanied by pulsed contractions is faster than constriction of boundary LECs without actin foci (Fig. 4G).

### Control of cellular contractility is crucial for enabling behavioural change

Overall, our results highlight the importance of contractility levels mediated by the amount of activated myosin, not only for the contractile network, but also for cellular architecture and cell behaviour (Fig. 8E). Contractility needs to be tightly controlled, otherwise LECs will change their behaviour. For wild-type LEC migration and the subsequent behavioural transition to constriction, a polarised cytoskeletal network and intermediate contractility levels seem to be crucial. However, rhythmical cytoskeletal activity appears not to be limited to rhythmical contractions of an actomyosin network, but also to extend to more general dynamic and rhythmical remodeling of the cytoskeletal architecture.

## Materials and methods

### Fly Stocks

FlyBase (Gramates et al., 2016) entries of the used transgenes are as follows: *hh.Gal4*: *Scer/Gal4^hh-Gal4^*, *UAS.gma-GFP*: *Moe^Scer\UAS.T:Avic\GFP-S65T^*, *UAS.LifeActin-Ruby*: *Scer\ABP140^Scer\UAS.T:Disc\RFP-Ruby^*, *sqh::GFP*: *sqh^RLC.T:Avic\GFP-S65T^*, *sqh^-/-^*: *sqh^Ax3^*, *UAS.MbsN300*: *Mbs^N300.Scer\UAS^*, *UAS.rok-RNAi*: *Rok^KK107802^* (VDRC 104675 (Dietzl et al., 2007)), *UAS.rok-CAT*: *Rok^CAT.Scer\UAS^*, *UAS.rho1-CA*: *Rho1^V14.Scer\UAS^*, *UAS.rho1*: *Rho1^UAS.cMa^*, *tub-FRT-CD2-FRT-Gal4*: *Rnor\CD2^A902^*, *UAS.mCD8-GFP*: *Mmus/Cd8a^Scer\UAS.T:Avic/GFP^*, *UAS.FLP*: *FLP1^Scer\UAS.cDa^*, *tub.Gal80ts*: *Scer\Gal80^ts.αTub84B^*, hs.FLP: *FLP1^hs.PS^*, Rok::GFP: *sqh.GFP-Rok*.

For the sqh::GFP experiments, we used *sqh[Ax3]; sqh::GFP; sqh::GFP* flies. Flies with one copy of sqh::GFP (*sqh[Ax3]/ w; + / +; sqh::GFP*/ +) showed similar labelling (Fig. S2C). The Rok-RNAi line has been used before (Rousso et al., 2015).

### Expression of transgenes in LECs

To express *UAS*-transgenes in the posterior compartment, *hh.Gal4* or *en.Gal4* was used. If transgenes were lethal when expressed throughout development, we used *tub.Gal80ts* to repress expression until 30-50h before imaging (*UAS.rok-RNAi; UAS.MbsN300, UAS.rok-CAT, UAS.rho1-CA, UAS.rho1*). Repression was released by shifting flies to the restrictive temperature (29°C). The behaviour of the cytoskeleton was not altered by Gal80ts presence; foci were localised in the back of migrating and in the centre of constricting LECs (Fig. S5B). For the *UAS.rok-CAT* experiments, we used different temperatures to release transgene repression. At the restrictive temperature, 35.3% of pupae showed a weak phenotype (n=17). At 25°C, where repression of Gal4 by Gal80ts is leaky, 100% of pupae showed a weak phenotype (n=5). All *UAS.rho1* pupae were grown at 25°C.

### 4D microscopy

For microscopy, pupae were staged according to (Bainbridge and Bownes, 1981). Pupae were dissected and filmed as described in (Seijo-Barandiarán et al., 2015). In all images and movies, anterior is to the left. All flies developed into pharate adults and many hatched. *Z*-stacks with a step size of 0.5 to 2.5 μm were recorded every 2 to 150s, depending on the experiment. Imaging was done with a Leica SP8 confocal microscope at 25±1°C. All images and movies shown are projections of *z*-stacks, if not mentioned otherwise. Figures and movies were made using Adobe Illustrator, Adobe Photoshop, Adobe Media Encoder, ImageJ (NIH, Bethesda, USA) and Leica LAS AF Lite (Leica Microsystems, Mannheim, Germany).

### Quantitative analysis of 4D movies

All LECs showed pulsed contractions and migratory and constrictive behaviour. We focussed our analysis on the dorsal side of segment A2, in a region at the back of the P compartment, three LEC rows away from the histoblasts at 18 h APF. This allowed comparability of individual cells, as cell shape, cell size and timing of delamination vary in different regions of the tissue.

#### a) Tracking of actin foci

Actin foci were tracked manually every 30s using the software SIMI Biocell (SIMI, Unterschleissheim, Germany) (Schnabel et al., 1997). Actin foci were defined as a local accumulation of fluorescence, in which fluorescence coalesced from different directions to then disassemble and that was visible for at least 3 frames. Movies for tracking were recorded over night with a time interval of 30s and a *z*-interval of 1 μm.

#### b) Calculating the relative position of actin foci

In addition to actin foci, the most anterior, posterior, dorsal and ventral coordinates of LECs were also tracked manually. These coordinates were used to calculate the relative position (RP) of foci in the cells along the a-p and the d-v axis (equations 1,2).

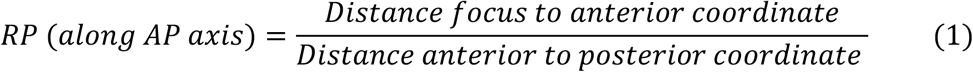

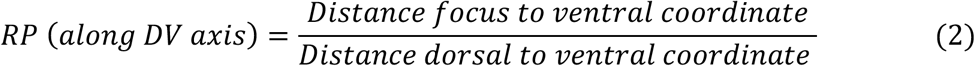

#### c) Calculating periodicity of actin foci

Actin foci were tracked and the time difference between two consecutive foci was calculated.

#### d) Calculating of cell shape coefficient

To obtain a single value that describes the shape of the cell, the cell shape coefficient was calculated using the most anterior, posterior, dorsal and ventral coordinates (equation 3).

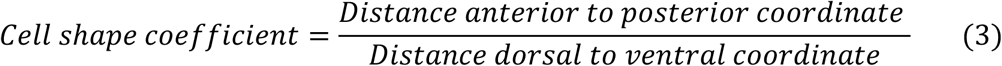

This coefficient assumes values close to 1 when the cell is round, and close to 0 when the cell is thin and long along the d-v axis.

#### e) Measuring cell area over time

Cell area was tracked manually using ImageJ. The polygon selection tool was used to draw the cell area, using as many vertices as needed to have an accurate outline. From one frame to the next, this selection was adjusted if cell area changed over time. To estimate the error associated with this technique, we tracked a cell for three frames and repeated the tracking 22 times. We found that the average error between repetitions was 0.37±0.07%. For seven wild-type LECs, cell area was measured every frame (30s) for the entire length of the recording, covering all four behavioural phases (Figs. 2; 3), For the analysis of mutant LECs, we measured a 30 min window of migratory and constrictive behaviour (beginning 25 min before lamellipodium disappearance (migration phase) and 75 min after lamellipodium disappearance (constriction phase)) (Figs. 5C; 6C; 7C). This allowed higher throughput. For the tracking of Rho-CA and Rok-CAT LECs we considered cell area excluding blebs.

#### f) Calculating reduction in cell area per contraction

The absolute magnitude of the cell area fluctuations was calculated by manually identifying the crests and the troughs for each fluctuation cycle and calculating the difference (scheme in Fig. 2B). Then the percentage by which the cell area is reduced in each fluctuation was calculated (equation 4).

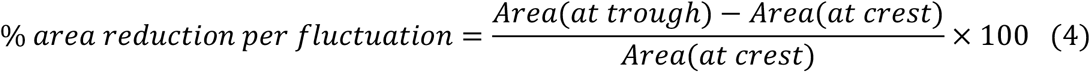

Differences in cell area reduction per contraction between the different phases of LEC behaviour are not due to differences in cell size, as similar results are obtained when considering percent area reduction and area reduction (in μm^2^) (compare Fig. 2B with Fig. S5C and Fig. 5C with Fig. S5D).

#### g) Assessing the number of actin foci per area fluctuation and the relationship between foci and area fluctuation

The number of actin foci per area fluctuation was calculated by manually counting the presence of foci tracked between the crests of each area fluctuation. To assess the relationship between actin foci and area fluctuation, we measured the time interval between a focus and the nearest trough of the area fluctuation by subtracting the time point at which the actin focus accumulates and the time point of the nearest trough for each tracked focus (scheme in Fig. 3C).

#### h) Co-localisation analysis of LifeActin-Ruby, Sqh::GFP and Rok::GFP

Using ImageJ, a region of 20 μm^2^ was drawn in the cell centre. Relative fluorescence intensities in this region were calculated for each channel using the plot profile function (Fig. 4A,B). This function creates a *y*-projection of the selected region before measuring intensities along the resulting line.

#### i) Cell size determination

As cells change shape dynamically and no two cells are in exactly the same stage of development at the same time, we needed to make sure that cell sizes could still be compared. For this, we measured the largest P compartment cell during migration and/or constriction using the polygon tool of ImageJ (Fig. 5B).

#### j) Kymographs

Kymographs were used to present cytoskeletal dynamics in a single image (Figs. 5E; 6B,E; 7A,B; 8A”,B). Kymographs were created from a rectangular region of interest (ROI), using the ImageJ tool ‘Reslice’, followed by a maximum *z*-projection of the obtained stack. This operation leads to vertical pixel rows, each of which depicts a y-projection of the ROI for each frame of the analysed movie (scheme in Fig. 5E). These rows were stacked on top of each other to illustrate the relative fluorescence intensity change in the ROI over time.

#### k) Trajectory plots

Trajectory plots (Fig. S3) were created using SIMI Biocell (Schnabel et al., 1997).

#### l) Quantification of abdominal closure timing and abdominal closure defects

For quantifying abdominal closure defects and assessing closure timing, *UAS*-transgenes were expressed in all LECs, as previously described (Bischoff, 2012). In short, FLP-out clones (Struhl and Basler, 1993) were induced by heat-shocking third-instar larvae for 15 min at 37°C. This led to transgene expression in all LECs but rarely in the histoblasts, as LECs are polyploid (Ninov et al., 2007). Expression of *UAS.FLP* increased recombination events in the polyploid LECs, thus increasing expression. After heat-shock, flies were kept at 25°C for 1-2 days before imaging.

To quantify how long it takes for LECs to complete abdominal closure (Fig. 5F’), we measured the time from the end of posterior migration (i.e. disappearance of posteriorly directed lamellipodia and start of histoblast nest expansion) to delamination of the last LEC.

#### m) Quantification of the duration of states of apicomedial and junctional cortical actin localisation in UAS.Rho1 pupae

To quantify cycling of LECs between the two phases (Fig. 8A”,C), fluorescence intensity was measured in a 23 μm^2^ area in a central apical region of the cells over time using ImageJ (see plot in Fig. 8A”). The resulting data were used to manually identify the two phases (‘high’ and ‘low’ fluorescence), as well as the transitions between the two phases, which were defined by a sudden drop/increase in fluorescence intensity with intensity values that lay outwith the range of intensities found during the ‘high’ or ‘low’ phases.

### Animal models and statistical analysis

statistical analyses were carried out in R (R core team, 2016) and Prism 8 (GraphPad Software, Inc., USA), assuming a significance level of 0.05. Sample sizes were determined by the available material that could be processed during each experiment. N-numbers for all experiments are shown in Table S1. No samples were excluded. To determine the appropriate statistical tests, the datasets were tested for normal distribution and for homogeneity of variance. Statistical tests for all experiments are shown in Table S2. For normally distributed data, Two-sample Student’s t-tests were used to compare the means of two groups. For more than two groups, one-way analyses of variance tests (ANOVA) were conducted. For non-parametric data of more than two groups, Kruskal-Wallis H tests were used, followed by a pairwise Wilcoxon-Mann-Whitney test to compare individual pairs. Medians were used for the non-parametric data, as they are better measurements of the central tendency of the data for skewed distributions. In order to calculate the standard errors and confidence intervals for the medians, a bootstrap method was applied (Efron, 1979), using a plugin in R. The number of replications chosen to obtain an accurate estimate was 1000 (used for analyses in Figs. 1E; 2B,C; 3E; 4H; 5C).

## Acknowledgements

We thank Verena Dietrich-Bischoff and Helder Ferreira for critically reading the manuscript and helpful discussions; Stefania Pasare, Malcolm White and Barry Denholm for critically reading the manuscript; Carl Robert Donavan for kindly providing the bootstrap code for R; Claudia Faustino for help with bootstrap code. Stocks obtained from the Bloomington *Drosophila* Stock Center (NIH P40OD018537) were used in this study. This work was supported by the BBSRC (BB/M021084/1).

## Author contributions

P.P.C. performed experiments, analysed data and co-wrote the manuscript. A.N. performed experiments, analysed data and commented on the manuscript. M.B. conceived the project, performed experiments, analysed data and wrote the manuscript.

## Notes

#### Summary of Updates

Supplementary files have been added to the initially submitted manuscript.

## References

Abreu-Blanco, M.T., J.M. Verboon, and S.M. Parkhurst. 2014. Coordination of Rho Family GTPase Activities to Orchestrate Cytoskeleton Responses during Cell Wound Repair. Curr Biol. 24:144–155.

Baena-Lopez, L.A., A. Baonza, and A. Garcia-Bellido. 2005. The orientation of cell divisions determines the shape of *Drosophila* organs. Curr Biol. 15:1640–1644.

Bainbridge, S., and M. Bownes. 1981. Staging the metamorphosis of *Drosophila* melanogaster. J Embryol Exp Morphol. 66:57.

Banerjee, D.S., A. Munjal, T. Lecuit, and M. Rao. 2017. Actomyosin pulsation and flows in an active elastomer with turnover and network remodeling. Nat Commun. 8:1121.

Barrett, K., M. Leptin, and J. Settleman. 1997. The Rho GTPase and a putative RhoGEF mediate a signaling pathway for the cell shape changes in *Drosophila* gastrulation. Cell. 91:905–915.

Bertet, C., L. Sulak, and T. Lecuit. 2004. Myosin-dependent junction remodelling controls planar cell intercalation and axis elongation. Nature. 429:667–671.

Bischoff, M. 2012. Lamellipodia-based migrations of larval epithelial cells are required for normal closure of the adult epidermis of *Drosophila*. Dev Biol. 363:179–190.

Bischoff, M., and Z. Cseresnyes. 2009. Cell rearrangements, cell divisions and cell death in a migrating epithelial sheet in the abdomen of *Drosophila*. Development. 136:2403–2411.

Blanchard, G.B., S. Murugesu, R.J. Adams, A. Martinez-Arias, and N. Gorfinkiel. 2010. Cytoskeletal dynamics and supracellular organisation of cell shape fluctuations during dorsal closure. Development. 137:2743–2752.

Bloor, J.W., and D.P. Kiehart. 2001. zipper Nonmuscle myosin-II functions downstream of PS2 integrin in *Drosophila* myogenesis and is necessary for myofibril formation. Dev Biol. 239:215–228.

Bosveld, F., I. Bonnet, B. Guirao, S. Tlili, Z. Wang, A. Petitalot, R. Marchand, P.-L. Bardet, P. Marcq, F. Graner, and Y. Bellaïche. 2012. Mechanical control of morphogenesis by Fat/Dachsous/Four-jointed planar cell polarity pathway. Science. 336:724–727.

Butler, L.C., G.B. Blanchard, A.J. Kabla, N.J. Lawrence, D.P. Welchman, L. Mahadevan, R.J. Adams, and B. Sanson. 2009. Cell shape changes indicate a role for extrinsic tensile forces in *Drosophila* germ-band extension. Nat Cell Biol. 11:859–864.

Cao, X., T. Kaneko, J.S. Li, A.D. Liu, C. Voss, and S.S. Li. 2015. A phosphorylation switch controls the spatiotemporal activation of Rho GTPases in directional cell migration. Nat Commun. 6:7721.

Carmona-Fontaine, C., H.K. Matthews, S. Kuriyama, M. Moreno, G.A. Dunn, M. Parsons, C.D. Stern, and R. Mayor. 2008. Contact inhibition of locomotion in vivo controls neural crest directional migration. Nature. 456:957–961.

Christodoulou, N., and P.A. Skourides. 2015. Cell-Autonomous Ca2+ Flashes Elicit Pulsed Contractions of an Apical Actin Network to Drive Apical Constriction during Neural Tube Closure. Cell Rep. 13:2189–2202.

Chrzanowska-Wodnicka, M., and K. Burridge. 1996. Rho-stimulated contractility drives the formation of stress fibers and focal adhesions. J Cell Biol. 133:1403–1415.

Coravos, J.S., F.M. Mason, and A.C. Martin. 2017. Actomyosin Pulsing in Tissue Integrity Maintenance during Morphogenesis. Trends Cell Biol. 27:276–283.

Dawes-Hoang, R.E. 2005. Folded Gastrulation, Cell Shape Change and the Control of Myosin Localization. Development. 132:4165–4178.

Dietzl, G., D. Chen, F. Schnorrer, K.C. Su, Y. Barinova, M. Fellner, B. Gasser, K. Kinsey, S. Oppel, S. Scheiblauer, A. Couto, V. Marra, K. Keleman, and B.J. Dickson. 2007. A genome-wide transgenic RNAi library for conditional gene inactivation in Drosophila. Nature. 448:151–156.

Efron, B. 1979. Bootstrap Methods: Another Look at the Jackknife. The Annals of Statistics. 7:1–26.

Fanto, M., U. Weber, D.I. Strutt, and M. Mlodzik. 2000. Nuclear signaling by Rac and Rho GTPases is required in the establishment of epithelial planar polarity in the *Drosophila* eye. Curr Biol. 10:979–988.

Fischer, S.C., G.B. Blanchard, J. Duque, R.J. Adams, A.M. Arias, S.D. Guest, and N. Gorfinkiel. 2014. Contractile and Mechanical Properties of Epithelia with Perturbed Actomyosin Dynamics. PLoS ONE. 9:e95695.

Gong, Y., C. Mo, and S.E. Fraser. 2004. Planar cell polarity signalling controls cell division orientation during zebrafish gastrulation. Nature. 430:689–693.

Graessl, M., J. Koch, A. Calderon, D. Kamps, S. Banerjee, T. Mazel, N. Schulze, J.K. Jungkurth, R. Patwardhan, D. Solouk, N. Hampe, B. Hoffmann, L. Dehmelt, and P. Nalbant. 2017. An excitable Rho GTPase signaling network generates dynamic subcellular contraction patterns. J Cell Biol. 216:4271–4285.

Gramates, L.S., S.J. Marygold, G.d. Santos, J.-M. Urbano, G. Antonazzo, B.B. Matthews, A.J. Rey, C.J. Tabone, M.A. Crosby, D.B. Emmert, K. Falls, J.L. Goodman, Y. Hu, L. Ponting, A.J. Schroeder, V.B. Strelets, J. Thurmond, P. Zhou, A. H., M. S.J., M. S.J., H. R.A., dos S. G., R. S., G. B.R., H. R.A., M. B.B., C. M.A., N.-J. S., N.-J. S., S. A.C., V. K.J.T., C. M., O.-S. D., K. W.A., M. G.H., B. S.M., F. R.D., V. K.J., K. E.Z., H. Y., K. E.V., A. J.S., P. R.A., E. J.T., M. A., D. G., B. P., A. B., B. E., W. J., B. S., C. L., J. G., R. M., and Z. R.P. 2016. FlyBase at 25: looking to the future. Nucleic Acids Res. 44:gkw1016.

Hatan, M., V. Shinder, D. Israeli, F. Schnorrer, and T. Volk. 2011. The *Drosophila* blood brain barrier is maintained by GPCR-dependent dynamic actin structures. J Cell Biol. 192:307–319.

He, L., X. Wang, H.L. Tang, and D.J. Montell. 2010. Tissue elongation requires oscillating contractions of a basal actomyosin network. Nat Cell Biol. 12:1133–1142.

Huang, C.-H., M. Tang, C. Shi, P.A. Iglesias, and P.N. Devreotes. 2013. An excitable signal integrator couples to an idling cytoskeletal oscillator to drive cell migration. Nat Cell Biol. 15:1–13.

Irvine, K.D., and E. Wieschaus. 1994. Cell intercalation during *Drosophila* germband extension and its regulation by pair-rule segmentation genes. Development. 120:827–841.

Keller, R. 2002. Shaping the vertebrate body plan by polarized embryonic cell movements. Science. 298:1950–1954.

Kinoshita, N., N. Sasai, K. Misaki, and S. Yonemura. 2008. Apical accumulation of Rho in the neural plate is important for neural plate cell shape change and neural tube formation. Mol Biol Cell. 19:2289–2299.

Lauffenburger, D., and A. Horwitz. 1996. Cell migration: a physically integrated molecular process. Cell. 84:359–370.

Lecuit, T., and L. Le Goff. 2007. Orchestrating size and shape during morphogenesis. Nature. 450:189–192.

Lee, A., and J. Treisman. 2004. Excessive Myosin Activity in Mbs Mutants Causes Photoreceptor Movement Out of the Drosophila Eye Disc Epithelium. Mol Biol Cell. 15:3285–3295.

Levayer, R., and T. Lecuit. 2012. Biomechanical regulation of contractility: spatial control and dynamics. Trends Cell Biol. 22:61–81.

Madhavan, M.M., and K. Madhavan. 1980. Morphogenesis of the epidermis of adult abdomen of *Drosophila*. J Embryol Exp Morphol. 60:1–31.

Mao, Y., A.L. Tournier, P.A. Bates, J.E. Gale, N. Tapon, and B.J. Thompson. 2011. Planar polarization of the atypical myosin Dachs orients cell divisions in *Drosophila*. Genes Dev. 25:131–136.

Martin, A.C., M. Kaschube, and E.F. Wieschaus. 2009. Pulsed contractions of an actin-myosin network drive apical constriction. Nature. 457:495–499.

Mason, F.M., and A.C. Martin. 2011. Tuning cell shape change with contractile ratchets. Curr Opin Genet Dev. 21:671–679.

Mason, F.M., M. Tworoger, and A.C. Martin. 2013. Apical domain polarization localizes actin--myosin activity to drive ratchet-like apical constriction. Nat Cell Biol. 15:926–936.

Mason, F.M., S. Xie, C.G. Vasquez, M. Tworoger, and A.C. Martin. 2016. RhoA GTPase inhibition organizes contraction during epithelial morphogenesis. J Cell Biol. 214:603–617.

Montell, D.J. 1999. The genetics of cell migration in *Drosophila melanogaster* and *C. elegans* development. Development. 126:3035–3046.

Munjal, A., J.-M. Philippe, E. Munro, and T. Lecuit. 2015. A self-organized biomechanical network drives shape changes during tissue morphogenesis. Nature. 524:351–355.

Nikolaidou, K.K., and K. Barrett. 2004. A Rho GTPase signaling pathway is used reiteratively in epithelial folding and potentially selects the outcome of Rho activation. Curr Biol. 14:1822–1826.

Ninov, N., D.A. Chiarelli, and E. Martin-Blanco. 2007. Extrinsic and intrinsic mechanisms directing epithelial cell sheet replacement during *Drosophila* metamorphosis. Development. 134:367–379.

Ohta, Y., J.H. Hartwig, and T.P. Stossel. 2006. FilGAP, a Rho-and ROCK-regulated GAP for Rac binds filamin A to control actin remodelling. Nat Cell Biol. 8:803–814.

Paluch, E., C. Sykes, J. Prost, and M. Bornens. 2006. Dynamic modes of the cortical actomyosin gel during cell locomotion and division. Trends Cell Biol. 16:5–10.

Pellegrin, S., and H. Mellor. 2007. Actin stress fibres. J Cell Sci. 120:3491–3499.

Prokopenko, S.N., A. Brumby, L. O’Keefe, L. Prior, Y. He, R. Saint, and H.J. Bellen. 1999. A putative exchange factor for Rho1 GTPase is required for initiation of cytokinesis in Drosophila. Genes Dev. 13:2301–2314.

Rauzi, M., P.-F. Lenne, and T. Lecuit. 2010. Planar polarized actomyosin contractile flows control epithelial junction remodelling. Nature. 468:1110–1114.

Ridley, A.J. 2003. Cell Migration: Integrating Signals from Front to Back. Science. 302:1704–1709.

Roh-Johnson, M., G. Shemer, C.D. Higgins, J.H. McClellan, A.D. Werts, U.S. Tulu, L. Gao, E. Betzig, D.P. Kiehart, and B. Goldstein. 2012. Triggering a Cell Shape Change by Exploiting Preexisting Actomyosin Contractions. Science. 335:1232–1235.

Rousso, T., E.D. Schejter, and B.-Z. Shilo. 2015. Orchestrated content release from Drosophila glue-protein vesicles by a contractile actomyosin network. Nature. 18:181–190.

Royou, A., C. Field, J.C. Sisson, W. Sullivan, and, R. Karess. 2004. Reassessing the Role and Dynamics of Nonmuscle Myosin II during Furrow Formation in Early *Drosophila* Embryos. Mol Biol Cell. 15:3751–3737.

Sander, E.E., J.P. ten Klooster, S. van Delft, R.A. van der Kammen, and J.G. Collard. 1999. Rac downregulates Rho activity: reciprocal balance between both GTPases determines cellular morphology and migratory behavior. J Cell Biol. 147:1009–1022.

Saravanan, S., C. Meghana, and M. Narasimha. 2013. Local, cell-nonautonomous feedback regulation of myosin dynamics patterns transitions in cell behavior: a role for tension and geometry? Mol Biology Cell. 24:2350–2361.

Sawyer, J.M., J.R. Harrell, G. Shemer, J. Sullivan-Brown, M. Roh-Johnson, and B. Goldstein. 2010. Apical constriction: a cell shape change that can drive morphogenesis. Dev Biol. 341:5–19.

Schnabel, R., H. Hutter, D.G. Moerman, and H. Schnabel. 1997. Assessing normal embryogenesis in *C. elegans* using a 4D-microscope: variability of development and regional specification. Dev Biol. 184:234–265.

Seijo-Barandiarán, I., I. Guerrero, and M. Bischoff. 2015. In Vivo Imaging of Hedgehog Transport in *Drosophila* Epithelia. Methods Mol Biol. 1322:9–18.

Simoes, S., B. Denholm, D. Azevedo, S. Sotillos, P. Martin, H. Skaer, J.C. Hombria, and A. Jacinto. 2006. Compartmentalisation of Rho regulators directs cell invagination during tissue morphogenesis. Development. 133:4257–4267.

Simoes, S., Y. Oh, M.F.Z. Wang, R. Fernandez-Gonzalez, and U. Tepass. 2017. Myosin II promotes the anisotropic loss of the apical domain during *Drosophila* neuroblast ingression. J Cell Biol. 216:1387–1404.

Solon, J., A. Kaya-Copur, J. Colombelli, and D. Brunner. 2009. Pulsed forces timed by a ratchet-like mechanism drive directed tissue movement during dorsal closure. Cell. 137:1331–1342.

Struhl, G., and K. Basler. 1993. Organizing activity of wingless protein in *Drosophila*. Cell. 72:527–540.

R Core Team. 2016. A language and environment for statistical computing. R Foundation for Statistical Computing, Vienna, Austria.

Teng, X., L. Qin, R. Le Borgne, and Y. Toyama. 2017. Remodeling of adhesion and modulation of mechanical tensile forces during apoptosis in *Drosophila* epithelium. Development. 144:95–105.

Tree, D.R., J.M. Shulman, R. Rousset, M.P. Scott, D. Gubb, and J.D. Axelrod. 2002. Prickle mediates feedback amplification to generate asymmetric planar cell polarity signaling. Cell. 109:371–381.

Valencia-Expósito, A., I. Grosheva, D.G. Míguez, A. González-Reyes, and M.D. Martín-Bermudo. 2016. Myosin light-chain phosphatase regulates basal actomyosin oscillations during morphogenesis. Nat Comm. 7:10746.

van Impel, A., S. Schumacher, M. Draga, H.-M. Herz, J. Grosshans, and H.A.J. Müller. 2009. Regulation of the Rac GTPase pathway by the multifunctional Rho GEF Pebble is essential for mesoderm migration in the *Drosophila* gastrula. Development. 136:813–822.

Vasquez, C.G., M. Tworoger, and A.C. Martin. 2014. Dynamic myosin phosphorylation regulates contractile pulses and tissue integrity during epithelial morphogenesis. J Cell Biol. 206:435–450.

Verdier, V., Guang-Chao-Chen, and J. Settleman. 2006. Rho-kinase regulates tissue morphogenesis via non-muscle myosin and LIM-kinase during *Drosophila* development. BMC Dev Biol. 6:38.

Warner, S.J., and G.D. Longmore. 2009. Cdc42 antagonizes Rho1 activity at adherens junctions to limit epithelial cell apical tension. J Cell Sci. 187:119–133.

Wilson, C.A., M.A. Tsuchida, G.M. Allen, E.L. Barnhart, K.T. Applegate, P.T. Yam, L. Ji, K. Keren, G. Danuser, and J.A. Theriot. 2010. Myosin II contributes to cell-scale actin network treadmilling through network disassembly. Nature. 465:373–377.

Xie, S., and A.C. Martin. 2015. Intracellular signalling and intercellular coupling coordinate heterogeneous contractile events to facilitate tissue folding. Nat Comm. 6:1–13.

Zallen, J.A., and E. Wieschaus. 2004. Patterned gene expression directs bipolar planar polarity in *Drosophila*. Dev Cell. 6:343–355.

