## Supplementary information for "Cellular contractility coordinates cytoskeletal dynamics and cell behaviour during *Drosophila* abdominal morphogenesis"

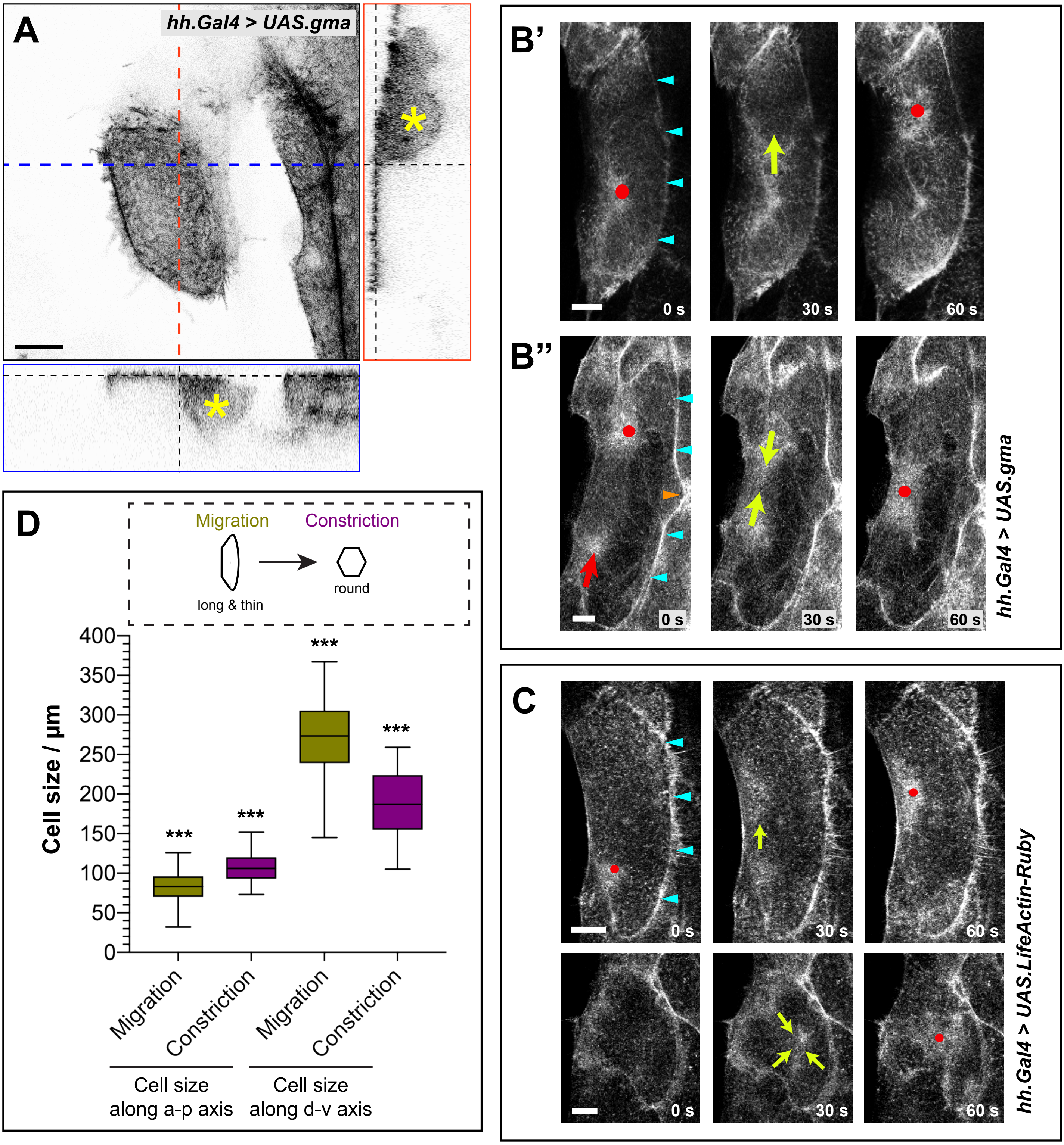
A
hh.Gal4 > UAS.gma
B'
0 s
30 s
60 s
B''
0 s
30 s
60 s
hh.Gal4 > UAS.gma
C
0 s
30 s
60 s
0 s
30 s
60 s
hh.Gal4 > UAS.LifeActin-Ruby
D
Migration
Constriction
long & thin
round
Cell size / µm
400
350
300
250
200
150
100
50
0
***
Migration
Constriction
Migration
Constriction
Cell size along a-p axis
Cell size along d-v axis

Figure S1

**Figure S1. The behaviour of the LEC actin cytoskeleton.** (**A**) LECs mainly consist of a large apical area with an apical actin network; F-actin labelled with GMA-GFP. The basal cell body surrounds the nucleus. Red, *z*-section along d-v axis. Blue, *z*-section along a-p axis. Asterisks indicate nucleus. (**B**) Actin flow patterns change when LECs transit from migration to constriction. GMA-GFP labels F-actin. Red dot, actin focus; yellow arrow, direction of actin flows; cyan arrowheads, lamellipodium. (**B'**) Typical pattern of actin foci and flows during migration – foci alternate between two positions and the flows move between these positions. (**B''**) During the transition from migration to constriction, disruption of lamellipodium shape (orange arrowhead) indicates the beginning of its disappearance. Actin flow patterns change: From the locations of the assembling (red dot) and disassembling (red arrow) foci, flows move towards the cell centre where the subsequent focus then forms. (**C**) Pulsed contractions and flows labelled with LifeActin-Ruby. Top, migration; bottom, constriction. Red dot, actin focus; yellow arrow, direction of actin flows; cyan arrowheads, lamellipodium. (**D**) Cell shape change during the transition from migration to constriction. Box plot showing the a-p and d-v length of LECs during migration and constriction. Cells are long and thin during migration and round during constriction. Most of the cell shape change is due to a shortening of the cells' d-v axes. All bars, 10 μm. Anterior, left; dorsal, up.

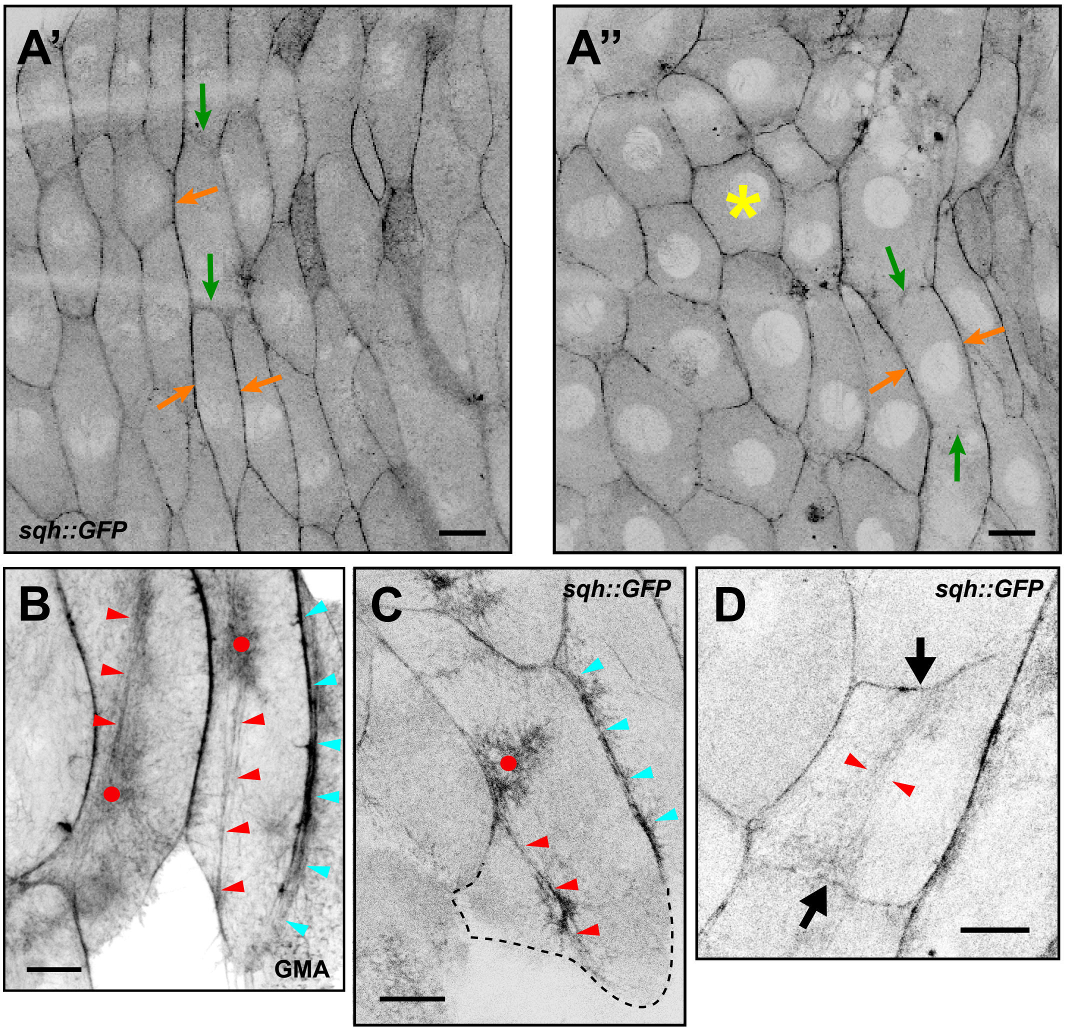


**Figure S2**

**Figure S2. Sqh::GFP localisation through morphogenesis.** (**A**) Overview of abdominal segment A2. (**A'**) At the beginning of morphogenesis, LECs preferentially localise Sqh::GFP at the a-p cell-cell interfaces (orange arrows), rather than the d-v interfaces (green arrows). (**A''**) During constriction (phase 3), cells that have not yet begun to constrict still localise Sqh::GFP preferentially at the a-p cell-cell interfaces (orange arrows), rather than the d-v interfaces (green arrows). Cells that are constricting have a more even Sqh::GFP localisation at all interfaces (asterisk). **(B)** Migrating GMA-GFP LECs showing actin foci (red dots) and actin bundles in the back (red arrowheads). Cyan arrowheads, lamellipodium. (**C**) Migrating LEC with two copies of Sqh::GFP shows focus (red dot) and actin bundles in the back (red arrowhead), as well as a lamellipodium at the front (cyan arrowheads). Dotted line indicates cell outline. (**D**) Constricting LECs occasionally show actin bundles that are oriented along the d-v axis (red arrowheads). These bundles pull at the dorsal and ventral membranes (black arrows). All bars, 10 µm. Anterior, left; dorsal, up.

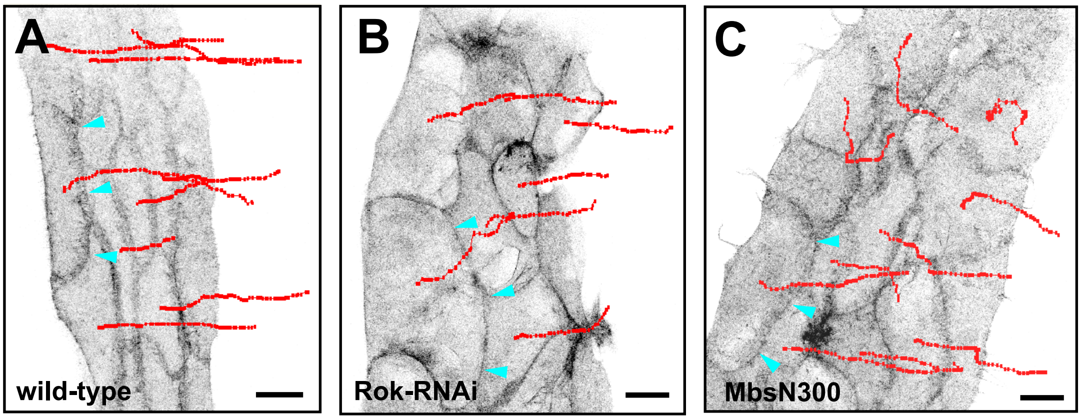


Figure S3

**Figure S3. Rok-RNAi and MbsN300 LECs migrate normally in posterior direction.** GMA-GFP labels F-actin. Tracks of cells shown in red; cyan arrowheads, lamellipodia. (**A**) GMA-GFP control. (**B**) Rok-RNAi. (**C**) MbsN300 overexpression. Bars, 20 μm. Anterior, left; dorsal, up.

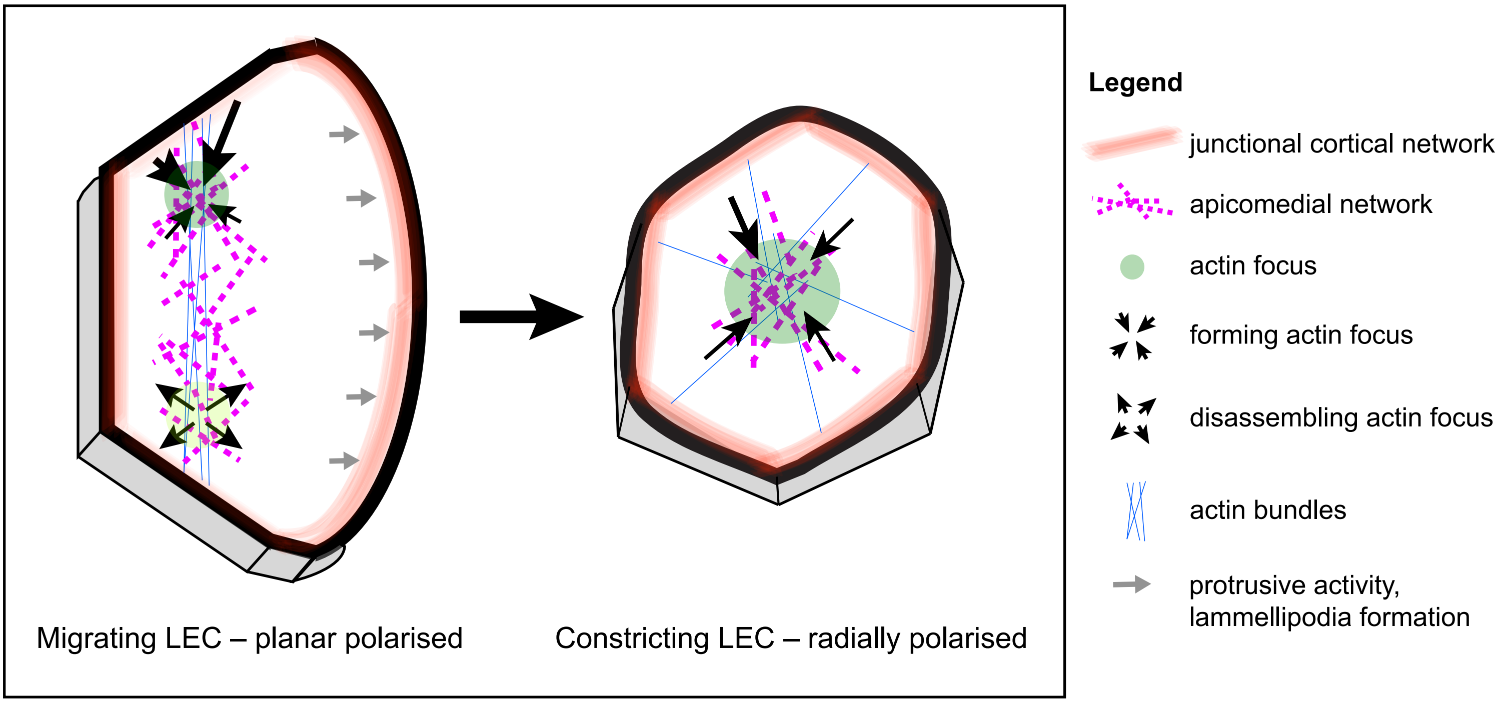
Legend
junctional cortical network
apicomedial network
actin focus
forming actin focus
disassembling actin focus
actin bundles
protrusive activity, lammellipodia formation
Migrating LEC – planar polarised
Constricting LEC – radially polarised

**Figure S4**

**Figure S4.** (**A**) **LECs undergo complex changes in cytoskeletal architecture during their behavioural switch from migration to constriction.** *Left*: A migrating LEC's cytoskeleton shows numerous features that indicate planar polarity. (a) Protrusive activity in the lamellipodium. (b) A contractile apicomedial network in the back of the cell. (c) Actin bundles that are oriented along the d-v axis in the back of the cell. (d) A planar polarised cortical network, which might contribute to contractility. (e) A basolateral contractile flow in the back of the cell. *Right*: A constricting LEC's cytoskeleton has a different architecture, which appears radially polarised. (a) A contractile medio-apical network in the centre of the cell. (b) Actin bundles that are oriented radially. (c) A cortical network, which localises around the whole cell and which might contribute to contractility. Anterior, left; dorsal, up.

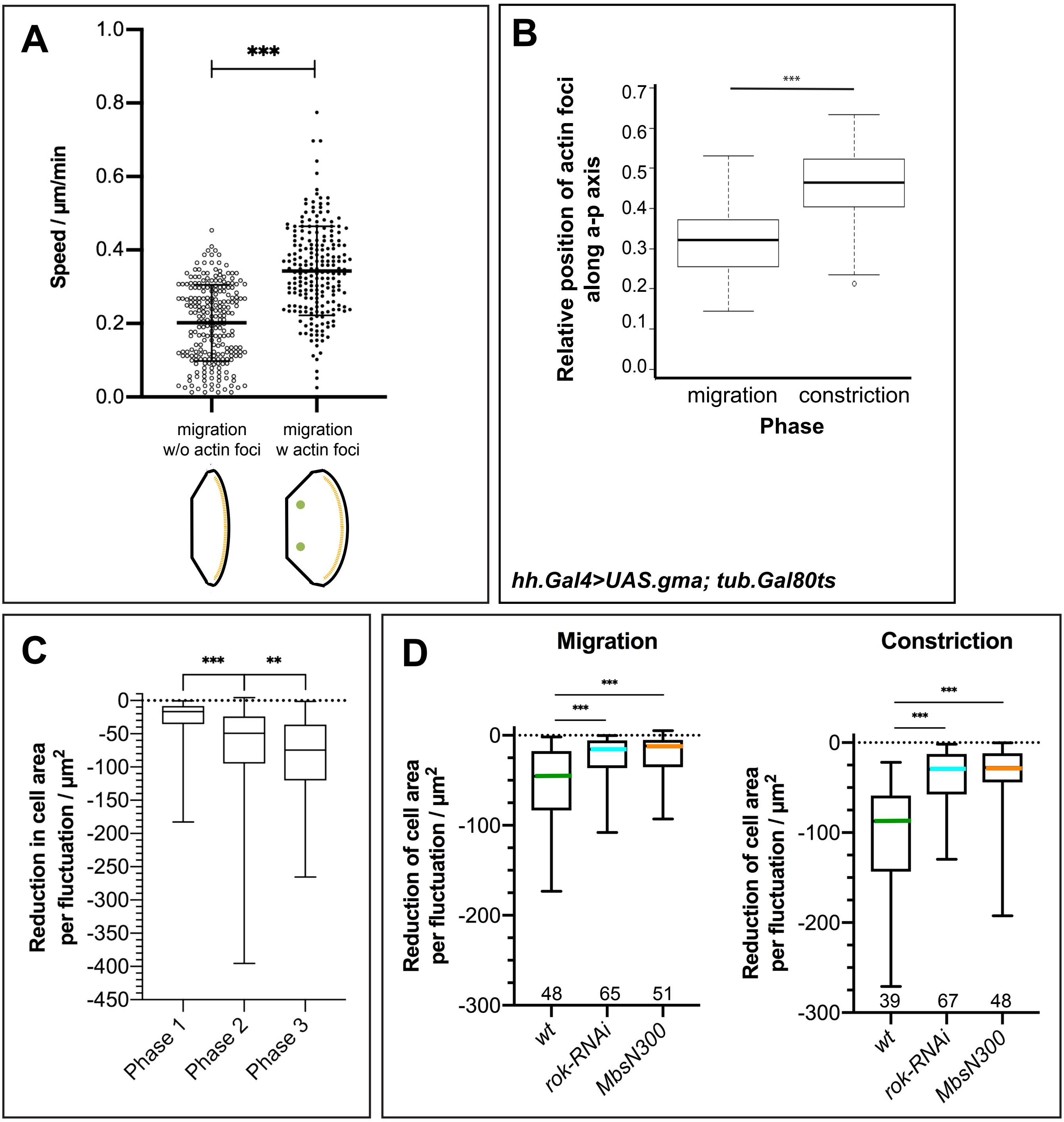
A
***
Speed / µm/min
1.0
0.8
0.6
0.4
0.2
0.0
migration w/o actin foci
migration w actin foci
B
***
Relative position of actin foci along a-p axis
0.7
0.6
0.5
0.4
0.3
0.2
0.1
0.0
migration
constriction
Phase
hh.Gal4>UAS.gma; tub.Gal80ts
C
***
**
Reduction in cell area per fluctuation / µm2
0
-50
-100
-150
-200
-250
-300
-350
-400
-450
Phase 1
Phase 2
Phase 3
D
Migration
Constriction
***
***
Reduction of cell area per fluctuation / µm2
0
-100
-200
-300
48
65
51
39
67
48
wt
rok-RNAi
MbsN300

**Figure S5**

**Figure S5.** (**A**) Comparing the speed of migration of LECs without actin foci (phase 1) and with actin foci (phase 2). LECs undergoing pulsed contractions migrate significantly faster. (**B**) Expression of Gal80ts does not alter positioning of actin foci. Boxplot showing relative position of actin foci along the a-p axis in LECs that express *UAS.gma-GFP* in a *tub.Gal80ts* background. In migrating cells, actin foci are found in the back, while in constricting cells, actin foci are located in the centre. The period between pulses is comparable to wild-type (180±0.2s; n= 5). (**C**) Boxplot comparing the reduction in cell area per pulsed contraction in phases 1, 2 and 3 (migration w/o actin foci, migration w actin foci, constriction; n=7). (**D**) Boxplot comparing reduction in cell area per pulsed contraction in wild-type, *rok-RNAi* and *UAS.MbsN300* during migration and constriction (*n* (foci) given along *x*-axis). **p<0.001, ***p<0.001.

### Table S1. Summary of *n*-numbers used.

| Experiment | *n*-number | Figure |
|---|---|---|
| *hh.Gal4 > UAS.gma-GFP* | | |
| - whole process (tracking) | 7[a] | 1C-F; 3A-C; 5A,D; S1B; S3A |
| - whole process (period) | 7[b] | (in text) |
| - whole process (area fluctuation) | 7 | 2A,B; 3A-C |
| - dorsal migration | 3[c] | 1G |
| - late phase 3 | 4 | 2C; 4H |
| - high magnification | 8 | 1B; 5C; S1A,D; S2C |
| - phase 2 and 3 (25/75min) | 4 | 5C; 6C; 7C |
| - cell area | 17 | 5B |
| - migration tracks | 7 | S3A |
| *hh.Gal4 > UAS.LifeActin-Ruby* | 7 | 1D; S1C |
| *hh.Gal4 > UAS.gma-GFP; tub.Gal80ts* | 5 | S1E |
| *sqh::GFP* | | |
| - tissue overview | 6 | S2A |
| - high magnification | 17 | 4C-E; S2B |
| - boundary LEC analysis | 4 | 4H |
| - Sqh::GFP *(one copy)* | 20 | S2D |
| *sqh[Ax3]; sqh-GFP/UAS.LifeActin-Ruby; sqh-GFP/+* | 14 | 4A |
| *sqh[Ax3]; rok::GFP/UAS.LifeActin-Ruby* | 20 | 4B |
| *UAS.mCD8-GFP* expression in all LECs to assess abdominal closure timing | 4 | 5F’ |
| *hh.Gal4 > UAS.rok-RNAi* | | |
| - weak phenotype | 6 | 5B,C,E |
| - strong phenotype | 2 | 5B,C,E |
| - high magnification | 11 | 5A |
| - migration tracks | 4 | S3B |
| - abdominal closure timing | 7 | 5F’ |
| *hh.Gal4 > UAS.MbsN300* | | |
| - weak phenotype | 1 | 5B,C,E |
| - strong phenotype | 4 | 5B,C,E |
| - high magnification | 13 | 5A |
| - migration tracks | 4 | S3C |

| | | |
|---|---|---|
| - abdominal closure timing | 6 | 5F’ |
| *hh.Gal4 > UAS.rok-CAT* | | |
| - weaker phenotype (25ºC) | 5 | 6A,B,C |
| - weaker phenotype (29ºC) | 6 | in text |
| - stronger phenotype (29ºC) | 11 | 6C,D,E |
| *hh.Gal4 > UAS.rho1-CA* | | |
| - weaker phenotype | 2 | 7A,C |
| - stronger phenotype | 5 | 7B,C |
| *hh.Gal4 > UAS.rho1* | 5[d] | 8 |

[a] sample size tested with Power test ($p<0.05$).

[b] 394 actin foci.

[c] 80 actin foci.

[d] 33 cells in 5 pupae

**Table S2. Statistical tests used.**

| Experiment | Figure | Test |
|---|---|---|
| - Relative position of actin foci<br>- Cell area | 1D,F; S5B<br>5B | ANOVA / t-test |
| - Period of actin foci<br>- Cell area fluctuations<br>- Change in cell shape<br>- Timing of abdominal closure | (in text)<br>2B,C; 3E; 4H; 5C; S5C,D<br>S1D<br>5F’ | Kruskal-Wallis H test |
| - Duration of phase<br>- Speed of migration | 8C<br>S5A | Mann-Whitney U test |

### Movie legends

**Movie S1. LECs undergo pulsed contractions that correlate with their behaviour.** Confocal micrographs of two LECs labelled with GMA-GFP undergoing pulsed contractions. Right cell is migrating, showing a lamellipodium (cyan arrowheads), and contracting at the same time. Left cell has lost its lamellipodium, has stopped migrating and is only constricting – orange arrowheads indicate the cell interface that is more flexible than the protruding lamellipodium of the migrating LEC. The migrating cell shows two alternating actin foci, the constricting cell shows one focus (red dots indicate foci). Red arrow, actin bundles in the back of the migrating cell. Bar, 10 µm. a, anterior; p, posterior; d, dorsal; v, ventral.

**Movie S2. Different phases of LEC behaviour. Confocal micrographs of LECs labelled with GMA-GFP.** During early migration (phase 1), LECs migrate without showing pulsed contractions. Merely flickering of apical activity is visible. During late migration (phase 2), LEC shows two actin foci alternating in their back (red dots). During constriction (phase 3), LEC shows an individual central actin focus (yellow dot). Cyan arrowheads, lamellipodium; hb, histoblasts; bars, 10 µm; a, anterior; p, posterior; d, dorsal; v, ventral.

**Movie S3. During dorsal repolarisation of a LEC, pulsatile activity reorganises to the back of the dorsally-migrating cell.** Confocal micrographs of LECs labelled with GMA-GFP. Before repolarisation, actin focus is in the cell centre (1st red dot). After repolarisation, actin focus is found in the back of the cell (2nd red dot). hb, histoblasts; bar, 10 µm; a, anterior; p, posterior; d, dorsal; v, ventral.

**Movie S4. Cytoskeletal architecture and pulsatile activity of a constricting LEC.** Sqh::GFP labels actin foci, actin bundles and cell-cell interfaces during constriction. Radially organised actin bundles (red arrowheads) connect the apicomedial network to the cell cortex. The contractile event begins in the cell periphery and then moves towards the cell centre. Red ellipsoids indicate movement of fluorescence signal towards the cell centre. Red dot shows actin focus at full contraction. Bar, 10 µm; a, anterior; p, posterior; d, dorsal; v, ventral.

**Movie S5. Constrictive behaviour of boundary LEC at the beginning of histoblast nest expansion, early during morphogenesis.** Sqh::GFP labels cell-cell interfaces. Also, some diffuse labelling in the apical cell area, but no actin foci are visible. LEC constricts over time. Bar, 10 µm; a, anterior; p, posterior; d, dorsal; v, ventral.

**Movie S6. Overview movie of Rok-RNAi LECs showing impaired contractile behaviour and actin flows in the back of the cells.** Confocal micrographs of LECs during late migration and constriction, F-actin labelled with GMA-GFP. LECs generate a lamellipodium in the front (cyan arrowheads) and show a more diffuse cytoskeleton labelling without actin foci (in area highlighted by yellow ellipsoid). Some cells create contractile flows in their back (cyan asterisks). LECs constrict and delaminate eventually (red arrow). Bar, 10 µm; a, anterior; p, posterior; d, dorsal; v, ventral.

**Movie S7. High magnification of two Rok-RNAi LECs showing impaired contractile behaviour and actin flows in the back of their neighbours.** Confocal micrographs of LECs during late migration, F-actin labelled with GMA-GFP. Cells generate lamellipodium in the front (cyan arrowheads) and show more diffuse actin foci compared to wild-type (Movie S1). The cell on the right and its right-hand neighbour create contractile flows in their back (cyan asterisks), which lie underneath the lamellipodium of their left-hand neighbour (orange line highlights the overlapping region of the cell on the right). Bar, 10 µm; a, anterior; p, posterior; d, dorsal; v, ventral.

**Movie S8. Overview of MbsN300 LECs showing impaired contractile behaviour and actin flows in their back.** Confocal micrographs of LECs during late migration and constriction, F-actin labelled with GMA-GFP. Cells generate lamellipodium in the front (cyan arrowheads) and show a more diffuse cytoskeleton labelling without actin foci (in area highlighted by yellow ellipsoid). Some cells create contractile flows in their back (cyan asterisks). LECs constrict and delaminate eventually (red arrow). Hb, histoblasts. Bar, 10 µm; a, anterior; p, posterior; d, dorsal; v, ventral.

**Movie S9. Rok-CAT overexpression in LECs at 25ºC, 'weak' phenotype.** F-actin labelled with GMA-GFP. Beginning of morphogenesis, migration and constriction visible. Constricting LECs show high levels of cortical actin (blue arrowheads). LECs migrate, showing lamellipodia (cyan arrowheads) and blebbing (magenta arrowheads). Some LECs show pulsatile behaviour (red dot). Bar, 10 µm; a, anterior; p, posterior; d, dorsal; v, ventral.

**Movie S10. Rok-CAT overexpression in LECs at 29ºC, 'strong' phenotype.** F-actin labelled with GMA-GFP. LECs are merely constricting, showing extensive blebbing (magenta arrowheads) and cortical actin bundles (orange arrowheads). No pulsatile behaviour visible. Eventually, cells delaminate (red arrow). Bar, 10 µm; a, anterior; p, posterior; d, dorsal; v, ventral.

**Movie S11. Rho1-CA overexpression in LECs, 'migrating' phenotype.** F-actin labelled with GMA-GFP. Beginning of morphogenesis, migration and constriction visible. LECs migrate, showing lamellipodia (cyan arrowheads); constricting LECs show blebbling (magenta arrowheads). No pulsatile behaviour visible. Bar, 10 µm; a, anterior; p, posterior; d, dorsal; v, ventral.

**Movie S12. Rho1-CA overexpression in LECs, 'non-migrating' phenotype.** Apical (left) and apicolateral (right) *z*-slice shown. F-actin labelled with GMA-GFP. LECs are merely constricting, showing extensive blebbing (magenta arrowheads). No pulsatile behaviour visible. Eventually, cells delaminate (yellow arrow). Bar, 10 µm; a, anterior; p, posterior; d, dorsal; v, ventral.

**Movie S13. Rho1 overexpression in LECs.** F-actin labelled with GMA-GFP. Orange asterisks indicate LECs that cycle between (1) the presence of apicomedial actin and (2) the absence of apicomedial actin but increased junctional cortical actin and blebbing. Pink arrowhead highlights LEC which is blebbing and shows junctional cortical actin. Bar, 20 µm. Anterior is to the left.
